# Pannexin 1 Ablation Disrupts Cell Cycle Progression and Compromises the Structure of Melanoma Tumours

**DOI:** 10.64898/2026.10.08.757604

**Authors:** Stephanie E. Leighton, Danielle Johnston, Kevin Barr, Brooke L. O’Donnell, Wyatt J. Schug, Matthew S. Huver, Dan Stefan, Christopher A. Bonham, John J Kelly, Ying Xia, Brant E. Isakson, John Ronald, Dale W. Laird, Silvia Penuela

## Abstract

Pannexin 1 (PANX1) is a channel-forming glycoprotein that has been implicated in regulating melanoma cell properties such as growth, migration, and invasion. Here we report that the CRISPR-Cas9 mediated ablation of *PANX1* or channel inhibition with probenecid shifted A375-P cell cycle distribution towards G2/M, while channel inhibition with spironolactone stalled cell cycle progression at the G1/S checkpoint regardless of PANX1 expression. Bulk RNA sequencing revealed cell cycle related gene expression was downregulated with PANX1 targeting strategies and this was exacerbated when *PANX1* deficient cells were treated with spironolactone. When grown as spheroids, we found the deletion and inhibition of PANX1 disrupted spheroid compaction, with spheroids displaying poorly defined borders and reduced circularity. The deletion of *PANX1* increased spheroid fragmentation, and this phenotype was intensified when spheroids devoid of PANX1 were treated with spironolactone, highlighting a potential synthetic lethal effect. Using liquid chromatography mass spectrometry, we sought to determine whether PANX1 had binding partners that may act to stabilize spheroid integrity and found that PANX1 co-immunoprecipitated various cell adhesion molecules such as β-actin and F-actin-capping protein subunit α-1, suggesting PANX1 and its interacting partners may stabilize cell-cell contacts. When control and PANX1 ablated cells were grown in chick chorioallantoic membrane xenografts to assess their tumour growth characteristics, we discovered that the deletion of *PANX1* reduced tumour surface area and viability. Our findings indicate that PANX1 regulates cell cycle progression and the structural architecture of A375-P melanoma tumours, highlighting the therapeutic utility of targeting PANX1 in melanoma.

## INTRODUCTION

Cutaneous melanoma is the deadliest form of skin cancer, and it arises from the malignant transformation of neural crest derived melanocytes that reside in the basal layer of the epidermis (1–3). Melanoma tumours progress through a radial growth phase where tumours proliferate horizontally followed by invasive vertical growth that leads to local invasion and metastasis (1,4,5). While surgical resection is the principal therapeutic approach for the successful treatment of primary melanoma, this intervention shows limited efficacy for metastatic disease (6). As such, the melanoma therapeutic landscape has evolved to include treatment options such as targeted therapies and immunotherapies, although approximately 60% of patients with metastatic disease do not respond to these treatment strategies (7–11). While recent therapeutic strategies show great promise in reducing the risk of recurrence in patients with high-risk resected melanoma, such as is the case with Moderna and Merck’s personalized mRNA vaccine intismeran, additional targets are essential for improving patient prognosis with metastatic disease where surgical resection and standard of care therapeutic approaches show limited efficacy (12).

Pannexins (PANX1, PANX2, and PANX3) are a family of integral membrane glycoproteins that canonically function to facilitate cellular signaling through the regulated passage of ions and metabolites up to 1 kDa in size (13–16). Among pannexins, PANX1 is the most ubiquitously expressed with reports of its presence in most mammalian tissue and cell types (17,18). As a result of its widespread expression, PANX1 has emerged as a key player in numerous physiological and pathological processes, including cancers such as melanoma (19–33). Conversely, PANX2 and PANX3 expression is more restrictive with no evidence of these pannexins being found in melanocytes (17,30). In patient melanoma tumours, PANX1 is highly expressed at all stages of disease metastasis, and its levels positively correlate with melanoma aggressiveness in isogenic mouse (B16-F0, B16-F10, B16-BL6) and human (A375-P, A375- MA2) melanoma cell lines (22,30,34).

PANX1 monomers are co-translationally inserted into the endoplasmic reticulum and subsequently N-link glycosylated (35,36). These high-mannose PANX1 glycoproteins oligomerize into heptamers where they are reported to function as calcium leak channels in the endoplasmic reticulum membrane (37,38). In addition to this intracellular channel function, PANX1 oligomers are transported to the Golgi apparatus for further glycosylation and ultimately trafficked to the plasma membrane where they canonically function as large pore channels (34,39–41). PANX1 channels are activated through various mechanisms including voltage, mechanical stimulation, caspase cleavage, purinergic signalling, and post-translational modifications where its function has been experimentally linked to key physiological processes such as apoptosis and inflammation (13,16,42–45). In the context of melanoma, inhibiting PANX1 channel activity using established PANX1 channel blockers probenecid (PBN) or carbenoxolone (CBX) *in vitro* using established melanoma cell lines and *ex vivo* via chick chorioallantoic membrane xenografts models significantly reduced melanoma cell growth and migration, suggesting that PANX1 channel activity may be important for the regulation of melanoma progression (22). Moreover, the inhibition of PANX1 with PBN or CBX reduced the proliferation and progression of high-risk neuroblastoma, thus supporting the therapeutic benefit of targeting PANX1 in a distinct neural crest derived cancer type (23). Mechanistically, PBN and CBX have been shown to inhibit PANX1 channels by binding to the extracellular loop region of PANX1 (46–48). A similar finding was reported in breast cancer, where the pharmacological inhibition of PANX1 channels with PBN or CRISPR/Cas9 mediated ablation of *PANX1* reduced cell invasion and reverted cells from an mesenchymal to epithelial phenotype (26). In addition, spironolactone (SPIR) has recently emerged as a PANX1 channel inhibitor following an unbiased small molecule screen (49). Although the exact mechanism of inhibition remains undefined, SPIR has been widely used to assess the channel dependent contributions of PANX1 primarily within the cardiovascular system and in the context of cancer (50–54). For example, in cutaneous squamous cell carcinoma, inhibiting PANX1 with either SPIR or PBN reduced cell growth and migration, suggesting that PANX1 is involved in promoting tumorigenic properties in a model of non-melanoma skin cancer (55). Similarly, reducing PANX1 protein with shRNA significantly reduced melanoma growth, migration, and increased melanin synthesis ultimately reverting melanoma cells to a melanocytic cell phenotype (22,30,34). Altogether these findings highlight PANX1 as a viable therapeutic target since blocking its expression or inhibiting channel function elicits a tumour suppressive effect. However, the exact mechanism governing the role of PANX1 in melanoma remains unclear.

In addition to PANX1’s role as a large pore channel protein, evidence has emerged implicating channel-independent functions of PANX1 in several key oncogenic signalling pathways. PANX1 has been linked to the Wnt/β-catenin signalling pathway as the carboxy-terminus of PANX1 directly binds the amino-terminus of the Wnt pathway effector β-catenin. This was shown to drive melanoma cell growth and mitochondrial metabolic activity (34). Similarly, in 3T3-L1 preadipocytes the deletion of *Panx1* significantly reduced the levels of total and non-phosphorylated β-catenin (56). In addition, studies have revealed PANX1 crosstalk with the Hippo/YAP/TAZ pathway in malignant melanoma, since PANX1 co-immunoprecipitated IQ motif-containing GTPase-activating protein 1 (IQGAP1)—a scaffold protein that directly binds β-catenin and YAP (57). YAP is a major transcriptional co-activator of the Hippo/YAP/TAZ pathway and is necessary for melanoma cells to transition between proliferative and invasive mesenchymal cell phenotypes (58). Moreover, the interaction between PANX1 and IQGAP1 has additionally been reported in mouse vascular smooth muscle cells (59). Cumulatively, these findings point to channel dependent and independent roles for PANX1 in melanoma; although further investigation is required to understand the interplay between these PANX1 functions and melanoma progression.

In this study, we investigated the role of PANX1 in controlling cellular properties associated with melanoma tumor growth in the human A375-P melanoma cell line where *PANX1* was ablated or inhibited with established PANX1 channel blockers. Our results demonstrate that the ablation of *PANX1* altered cell cycle-related gene expression, shifting cells towards Gap 2/ Mitosis (G2/M) which was phenocopied by the inhibition of PANX1 with PBN, highlighting a role for PANX1 in modulating cell cycle progression. We found that treatment SPIR stalled cell cycle progression at the Gap 1/ Synthesis (G1/S) checkpoint regardless of PANX1 expression highlighting pleotropic effects of SPIR in melanoma cells. Bulk RNA sequencing revealed that genes within the cell cycle pathways, E2F target and G2/M checkpoint were downregulated with PANX1 targeting strategies and this was exacerbated when *PANX1* deficient cells were treated with spironolactone. This finding ultimately reveals a previously undefined synergy between PANX1 expression and spironolactone. We also demonstrate that PANX1 is involved in promoting spheroid assembly and adhesivity since the deletion of *PANX1* reduced spheroid circularity and increased fragmentation of the primary spheroid following mechanical stress. Which likely occurs due to PANX1 interacting with various cell adhesion molecules such as β-actin and F-actin-capping protein subunit α1 that may act to stabilize the integrity of the spheroids. We further demonstrate that the deletion of *PANX1* reduced *en face* tumour area and viability when assessed *ex vivo* in a chick chorioallantoic xenograft model, emphasizing the therapeutic benefit of targeting PANX1 in melanoma.

## MATERIALS AND METHODS

### Cell Culture

Wild-type (WT) A375-P cells (RRID:CVCL_6233) purchased from the American Type Culture Collection (ATCC, Manassas, VA, USA), scrambled control (SC), and *PANX1* knock out (PANX1 KO) A375-P cells were incubated in sterile T75 flasks (Fisher Scientific, Cat# FB012937) at 37°C and 5% CO_2_ in Dulbecco’s Modified Eagle Medium (DMEM) (Thermo Fisher Scientific, Cat# 12430062) that was supplemented with 10% fetal bovine serum (Wisent Inc., Cat# 098-150) and 100 units/mL penicillin-streptomycin (Thermo Fisher Scientific, Cat# 15070063). Cells were washed once with calcium and magnesium free 1x Dulbecco’s phosphate buffered saline (PBS) (Thermo Fisher Scientific, Cat# 14190250) and subsequently dissociated with 0.25% trypsin/ 1 mM EDTA (Wisent Inc., Cat# 325-043-EL) for subculturing. Where noted, cells were counted using a trypan blue exclusion assay (Thermo Fisher Scientific, Cat# 15250061) with the Countess II FL (Thermo Fisher Scientific). The MycoAlert Mycoplasma Detection Kit (Lonza, Cat# LT07-318) was used to routinely test cells for mycoplasma contamination.

### CRISPR/Cas9 Gene Editing and Clonal Expansion

WT A375-P cells were nucleofected using the SF Cell Line 4D-Nucleofector® X Kit S (Lonza, Cat# V4XC- 2032), where ribonucleoprotein complexes were assembled by combining 1.2 µL of sgRNA (100 µM) (5’- GGCCAGUUGAGCGAUGGCCA-3’, 5’-GCCCACGGAGCCCAAGUUCA-3’, and 5’- GAUGGUCACGUGCAUUGCGG-3’ or negative control sgRNA 5’-GCACUACCAGAGCUAACUCA-3′) purchased from Synthego (Menlo Park, CA, USA) and 1 μL of HiFi Cas9 Nuclease (20 μM) at room temperature for 10 minutes. Two hundred thousand WT A375-P cells per reaction were resuspended in 20 μL of SF 4D-Nucleofector® X solution and supplement (4.5:1 ratio) and subsequently combined with the ribonucleoprotein complexes according to the manufacturer’s protocol (Lonza, Cat# V4XC-2031). The reaction mixtures were transferred into a 16-well Nucleocuvette® vessel and nucleofected using the 4D- Nucleofector® X Unit (Lonza, Cat# AAF-1003X) with the pre-set program for A375 cells. Following nucleofection, cells were incubated at room temperature for 5 minutes and subsequently resuspended in 80 μL of DMEM for a 10 minute recovery incubation at 37°C and 5% CO_2_. Cells were then seeded into a 24-well plate (Greiner Bio-One, Cat# 662165) and grown to confluence. Eight days following nucleofection, cells were subcultured into a 96-well plate (Greiner Bio-One, Cat# 655 180) for clonal expansion using limiting dilution. Single cell colonies were identified using the Olympus CK40 Culture Microscope and expanded for downstream analysis. Successful ablation of *PANX1* was validated via immunoblotting and immunofluorescence microscopy. Once clones were independently validated, a subset of SC (SC1, SC2, SC3, SC4) and PANX1 KO (KO1, KO2, KO3, KO4) A375-P clones were pooled at an equal cell number to form a mixed clonal population for downstream analysis.

### Monolayer Protein Extraction and Immunoblotting

WT, SC, and PANX1 KO A375-P cells were grown as a monolayer on 60 mm tissue culture dishes (Fisher Scientific, Cat# FB012921) until confluent for cell lyses on ice with 1x RIPA lysis buffer (50 mM Tris-HCl at pH 8.0,150 mM NaCl, 1% NP-40, 0.5% sodium deoxycholate, 0.1% sodium dodecyl sulphate) supplemented with 1 mM sodium fluoride (BDH, Cat #BDH9290-500G), 1 mM sodium orthovanadate (ThermoFisher Scientific, Cat #13721-39-6), and a complete™ mini protease inhibitor cocktail tablet (MilliporeSigma, Cat# 11836153001). In cases where SC and PANX1 KO A375-P cells were treated with PANX1 channel inhibitors PBN (1 mM; Invitrogen, Cat# P36400), or SPIR (10 µM; SelleckChem, Cat# S4054), three hundred and fifty thousand cells were seeded onto 60 mm tissue culture dishes and allowed to adhere for twenty-four hours before treating with either PBN or SPIR or their respective vehicle control (ultrapure water (Life Technologies, Cat# 10977-015) or 100% ethanol, respectively). Cell culture media supplemented with PANX1 inhibitors or vehicle was replaced forty-eight hours following the initial treatment and cells were grown for a total of five days. As a positive control for endoplasmic reticulum stress and apoptotic cell death, SC A375-P cells were treated with 2 μg/mL of tunicamycin (MilliporeSigma, Cat# 11089-65-9) for twenty-four hours or 1 μM staurosporine (MilliporeSigma, Cat# S6942) for three hours, respectively. Adherent cells were isolated and resolved by SDS-PAGE as previously described (60). The BLUelf Prestained Protein Ladder (FroggaBio, Cat# PM008) and prepared protein lysates were transferred to nitrocellulose membranes (Thermo Fisher Scientific, Cat# IB23001; IB23002; IB301002; or IB301001) using the iBlot or iBlot 2 dry-transfer system. Membranes were blocked with 3% bovine serum albumin (BSA) (BioShop, Cat# ALB001.100) in 0.05% Tween-20 phosphate buffered saline (PBS-T) and incubated with rabbit anti-PANX1 (CT-412) (0.35 µg/mL, Penuela et al., 2007(35)), rabbit anti-PANX1 (130 ng/mL, Cell Signaling Technology, Cat# 91137, RRID:AB_2800167), rabbit anti-GADD 153 (CHOP) (200 ng/mL, Santa Cruz Biotechnology, Cat# sc-575, RRID:AB_631365), rabbit anti-binding immunoglobulin protein (BiP) (6.1 µg/mL, Sigma-Aldrich Cat# G8918, RRID:AB_477030), rabbit anti-LC3B (1 µg/mL, Thermo Fisher Scientific Cat# PA1-46286, RRID:AB_2234770), rabbit anti-cleaved caspase 3 (cCAS3) (52 ng/mL, Cell Signaling Technology Cat# 9661, RRID:AB_2341188) primary antibodies overnight at 4°C or at room temperature for one hour in the case of mouse anti-GAPDH (0.04 ng/mL, Millipore, Cat# MAB374, RRID:AB_2107445). Primary antibody incubation was followed by goat anti-mouse IRDye® 800CW (50 ng/mL, LICORbio, Cat# 926-32210, RRID:AB_621842), goat anti-mouse IRDye® 680RD (50 ng/mL, LICORbio, Cat# 926-68070, RRID:AB_10956588), goat anti-rabbit IRDye® 800CW (50 ng/mL, (LICORbio, Cat# 926-32211, RRID:AB_621843), or goat anti-rabbit IRDye® 680RD (50 ng/mL, LICORbio, Cat# 926-68071, RRID:AB_10956166) secondary antibodies at room temperature for fourty-five minutes. Immunoblots were imaged using an Odyssey LiCor infrared imaging system and quantified with Odyssey Application Software (LI-COR).

### Monolayer Immunolabelling and Imaging

SC and PANX1 KO A375-P cells grown as a monolayer on 60 mm tissue culture dishes containing 12 mm glass coverslips (Fisher Scientific, Cat# S000845) were fixed and immunolabelled as described (61). To assess the consequences of PANX1 inhibition, three hundred and fifty thousand SC and PANX1 KO A375-P cells were seeded and left to adhere for twenty-four hours prior to treatment with 1 mM PBN, 10 µM SPIR, or vehicle controls. For all experiments, cell culture media containing the respective PANX1 inhibitor or control was replaced every other day up to five days. In all instances, cells were incubated overnight at 4°C with rabbit anti-PANX1 (CT-412) (0.7 µg/mL, Penuela et al., 2007(35)) followed by an Alexa Fluor 488 goat anti-rabbit secondary antibody (2.9 µg/mL, Thermo Fisher Scientific, Cat# A-11008, RRID:AB_143165) at room temperature for one hour. Nuclei were stained with Hoechst 33342 (1:1000; Thermo Fisher Scientific, Cat# H3570, RRID:AB_3675235) for five minutes at room temperature. Coverslips were mounted onto microscope slides (Fisherbrand, Cat #22034486) using prolong gold antifade mounting medium (Invitrogen, Cat # P36934). Confocal images were captured on a Carl Zeiss LSM 800 Airyscan confocal microscope using a plan-apochromat (63x, 1.4 NA) oil lens as described (62).

### Illumina Next Generation Sequencing

RNA was isolated from four biologically independent SC (SC1, SC2, SC3, SC4) and PANX1 KO (KO1, KO2, KO3, KO4) A375-P clones treated with 10 µM SPIR (SelleckChem, Cat# S4054) or vehicle (EtOH) for five days using the RNeasy mini kit (Qiagen, Cat#74104) as per manufacturer’s directions. All samples were sequenced at the London Regional Genomics Centre (Robarts Research Institute, London, Ontario, Canada; http://www.lrgc.ca) according to standard protocols using the Illumina NextSeq 500 (Illumina Inc., San Diego, CA) as described (63–65). Total RNA was quantified using the NanoDrop (Thermo Fisher Scientific, Waltham, MA) and quality was assessed using the Agilent 2100 Bioanalyzer (Agilent Technologies Inc., Palo Alto, CA) and the RNA 6000 Nano kit (Caliper Life Sciences, Mountain View, CA). Samples were processed using the Vazyme VAHTS V8 RNA-seq Library Prep Kit for Illumina (Vazyme, Nanjing, China). Libraries were then equimolar pooled into one library and size distribution was assessed on an Agilent High Sensitivity DNA Bioanalyzer chip and quantitated using the Qubit 2.0 Fluorimeter (Thermo Fisher Scientific, Waltham, MA). The library was sequenced on an Illumina NextSeq 500 as a 1 x 76 bp single end run, using a High Output v2 kit (75 cycles). Fastq data files were analyzed using Partek Flow (St. Louis, MO). After importation, data was aligned to the Homo sapiens genome using STAR 2.7.8a and annotated using hg38 Ensembl Transcripts release 110. Features with less than 5 reads were filtered out, followed by normalization using DESeq2 where fold change and p-values were determined (66). Filtered lists of genes changing 1.5-fold and with a p-value of less than 0.05 were then analyzed for enriched Kyoto Encyclopedia of Genes and Genomes (KEGG) pathway and Gene Ontology terms (67–69). Gene set enrichment analysis (GSEA) using the Molecular Signatures Database (MSigDB.v.2025.1.Hs) was performed using the normalized gene counts determined using DESeq2 (70,71).

### Adherent (2D) and Spheroid (3D) Cell Growth Assays

For adherent cell growth assays, SC1, SC2, SC3, SC4, SC5, SC6, PANX1 KO1, PANX1 KO2, PANX1 KO3, PANX1 KO4, PANX1 KO5, PANX1 KO6, PANX1 KO7, PANX1 KO8, PANX1 KO9, PANX1 KO10, PANX1 KO11, and PANX1 KO12 A375-P clones were seeded in triplicate at ten thousand cells/well in a 24-well plate (Greiner Bio-One, Cat# 662165). Then, using the Incucyte S3 Live-Cell Analysis Instrument (RRID:SCR_023147), phase contrast images were captured using the 10x objective every 4 hours to assess changes to cell growth over four to five days as described (55). In cases where SC1, SC2, SC3, SC4, PANX1 KO1, PANX1 KO2, PANX1 KO3, and PANX1 KO4 A375-P cells were challenged with PANX1 inhibitors, twenty-four hours after seeding cells were treated with 1 mM PBN, 10 µM SPIR, or vehicle control. Cell culture media supplemented with the corresponding treatment was replaced every other day. Cell counts were analyzed using the Incucyte Cell-By-Cell Analysis Software Module and triplicate measurements were averaged (55,72). The growth of each A375-P clone was assessed with three to four independent repeats, the sum of which was averaged for each clone.

For spheroid growth assays, six technical replicates of pooled SC or PANX1 KO A375-P cells were plated at two thousand cells/well in a 96-well ultra-low attachment round bottom plate (Costar, Cat# 7007) and allowed to spontaneously form spheroids for six days (Fisher Scientific, Cat# 07-200-601) (73–75). Phase contrast images were captured every four hours using the Incucyte S3 Live-Cell Analysis Instrument 10x objective. Spheroid area and circularity were analyzed using Fiji ImageJ Version 2.16.0/1.54p; Java 1.8.0_322[64-bit] for four independent replicates (76).

### Flow Cytometry

In a 6-well plate (Genesee Scientific, Cat# 25-105MP), twenty thousand SC or PANX1 KO A375-P cells were seeded per condition. Twenty-four hours after seeding, cell cycle synchronization was performed by incubating cells with 10% charcoal-stripped FBS (Thermo Fisher Scientific, Cat# A3382101) in DMEM for twenty-four hours. Cells were then treated with either 1 mM PBN, 15 µM SPIR or corresponding vehicle control in complete DMEM media (10% FBS, 1% Pen/Strep), and treatments were replaced again after two days. Six days after seeding, cells were washed once with 1x PBS, trypsinized with 1x TrypLE Express, no Phenol Red (Gibco, Cat# 12604021) for five minutes and counted using the Countess 3 FL (Thermo Fisher Scientific). Aliquots of seven hundred fifty thousand cells per condition were centrifuged at 800 g for 5 minutes at 4°C and pellets were resuspended in ice cold 70% ethanol for thirty minutes fixation on ice. After centrifugation for 10 minutes at 800 g at 4°C, cells were resuspended in 10 µM of DRAQ5 DNA stain (Thermo Fisher Scientific, Cat# 62254) diluted in 1x PBS. A375-P cells without treatment were resuspended in 1x PBS instead of DRAQ to use as an unstained control to determine gating strategies. Cells were covered from light, incubated for five minutes on ice and then analyzed on an Attune NxT Flow Cytometer (Thermo Fisher Scientific) using the following settings: flow rate 25 µL/min, forward scatter 110, side scatter 320, YL3 laser 430. Collected events were gated to exclude debris and doublets, and the same gating strategy was applied to all samples. Results were analyzed using FCS Express™ version 7 (De Novo Software) and G0/G1, S and G2/M cell cycle phases were manually segmented.

### Spheroid (3D) Cell Viability Assays

Two thousand SC or PANX1 KO A375-P cells per well were seeded in a 96-well ultra-low attachment round bottom plate. Immediately following seeding, spheroids were treated with 10 µM SPIR or vehicle control. Cell culture media supplemented with 10 µM SPIR or vehicle (EtOH) was replaced every fourty- eight hours. After a period of six days, spheroids were incubated for thirty minutes at 37°C in complete DMEM containing 5 µM Calcein AM (Invitrogen, Cat# C3100MP). Spheroids were subsequently washed twice with 1x PBS then imaged using the Incucyte S3 Live-Cell Analysis Instrument 4x objective. Calcein AM fluorescence intensity was analyzed using the Incucyte S3 Live-Cell Analysis Spheroid Analysis Software Module. Spheroid fragments were counted using the Fiji ImageJ Version 2.16.0/1.54p; Java 1.8.0_322[64-bit] cell counter plugin (76).

### Spheroid (3D) Immunolabelling and Imaging

For imaging whole spheroids, two thousand SC or PANX1 KO A375-P cells were seeded per well in a 96- well ultra-low attachment round bottom plate. After six days, spheroids were fixed at room temperature for thirty minutes with 4% paraformaldehyde (Thermo Fisher Scientific, Cat# 043368.9M). Fixed spheroids were then permeabilized at room temperature for fifteen minutes with 0.25% Triton X-100 (Amresco, Cat# 0694-1L) and subsequently blocked and immunolabelled as described above for adherent cells. Spheroids were immunolabelled with rabbit anti-PANX1 (CT-412) (1.75 µg/mL, Penuela et al., 2007(35)) or rabbit anti-PANX1 (1 µg/mL, Sigma-Aldrich, Cat# HPA016930, RRID:AB_1854954) primary antibodies and Alexa Fluor 488 goat anti-rabbit secondary antibody (2.9 µg/mL, Thermo Fisher Scientific, Cat# A- 11008, RRID:AB_143165). Whole spheroids were imaged on a Carl Zeiss LSM 800 Airyscan confocal microscope in the 96-well ultra-low attachment round bottom plate using a plan-neofluar 10x 0.3 numerical aperture.

### Adherent (2D) Cell Migration Assay

SC and PANX1 KO A375-P cells were grown to confluence in a Incucyte Imagelock 96-well plate (Sartorius, Cat# 4379) where a scratch wound was created in each well using the Incucyte Woundmaker Tool (Sartorius BA-04858) as described previously (55). After wounding, the cell monolayer was washed once with 1x PBS then grown in DMEM that was supplemented with 10% charcoal stripped fetal bovine serum (ThermoFisher, Cat#A33821-01), 100 units/mL penicillin-streptomycin, and the corresponding PANX1 inhibitor (1 mM PBN, 10 µM SPIR, or vehicle controls). Phase contrast images were captured every two hours using the Incucyte S3 Live-Cell Analysis Instrument to assess changes to cell migration over time. Overall wound closure was quantified using the Incucyte Scratch Wound Analysis Software Module.

### Immunoprecipitation

Once confluent, SC and PANX1 KO A375-P cells were lysed on ice with 1x immunoprecipitation buffer (10 mM Tris-HCl at pH 7.4, 150 mM NaCl, 1 mM EDTA, 1 mM EGTA, 0.5% NP-40 and 1% Triton X-100) supplemented with 1 mM NaF, 1 mM Na_3_VO_4_, and a cOmplete™ mini protease inhibitor cocktail tablet. Cell lysates were centrifuged at 4°C for fifteen minutes using the Hettich MIKRO 22 R, (Cat# 1110) centrifuge at 10,000 g. The resulting supernatants were collected for a bicinchoninic acid (BCA) assay (Thermo Fisher Scientific, Cat# 23277) to isolate 1 mg of protein per immunoprecipitation reaction. The lysates were diluted to a final volume of 250 µL with complete 1x immunoprecipitation buffer. Twenty-five µL (0.75 mg) of Pierce™ Protein G Magnetic Beads (ThermoFisher Scientific, Cat #88847) were isolated and washed twice with 0.05% PBS-T. The beads were subsequently combined with rabbit anti-PANX1 (CT-412) (10 μg/mL, Penuela et al., 2007(35)) that was diluted in 250 µL of 0.05% PBS-T and rotated for ten minutes at room temperature. The beads were then washed with 0.05% PBS-T three times and subsequently combined with 1 mg of protein lysate which was rotated for thirty minutes at room temperature and processed for mass spectrometry. In parallel, in order to validate the successful immunoprecipitation of PANX1, SC and PANX1 KO A375-P cells immunoprecipitated with rabbit anti- PANX1(CT-412) or rabbit anti-IgG (10 μg/mL; Thermo Fisher Scientific, Cat# 02-6102, RRID:AB_2532938) and eluted as described for validation via immunoblotting (60).

### Sample Processing for Mass Spectrometry

Processing of samples for mass spectrometry-based proteomic analysis was performed in conjunction with the Biomolecular Core (BioCORE) Facility at Western University Schulich School of Medicine and Dentistry, London, Canada. Briefly, magnetic affinity supports were washed with 0.05% PBS-T twice, followed by three washes with freshly prepared 50 mM NH_4_HCO_3_ pH 8 (Fisher Scientific) while transferring to fresh siliconized microcentrifuge tubes. Magnetic affinity supports were washed twice more with 50 mM NH_4_HCO_3_ pH 8. After the final wash, affinity supports were resuspended as a ∼20% slurry with 50 mM NH_4_HCO_3_ pH 8, then bound proteins were reduced with 5 mM Tris(2-carboxyethyl)phosphine (TCEP; ThermoFisher Scientific) and alkylated with 10 mM iodoacetamide (Sigma) with 1000 rpm shaking at 30°C for approximately fourty-fiveminutes. The bound protein was then digested with 1 μg sequencing grade lysyl endopeptidase (LysC; FUJIFILM Wako Chemicals) by shaking at 1400 rpm for four hours at 37°C, followed by the addition of 1 μg trypsin (MS grade; ThermoFisher Scientific) and left for overnight digestion as above. Sample peptide supernatants were collected, then beads were washed twice with 2% acetonitrile (ACN) containing 0.1% trifluoroacetic acid (TFA). Washes were pooled with peptide digests, and digests were further acidified to 0.5% TFA. Peptides were desalted using Oasis HLB 1cc (10 mg; Waters) solid-phase extraction cartridges, eluates evaporated to dryness by speed vacuum concentration, and resultant peptides were resuspended in 2% ACN, 0.1% TFA by shaking at 1400 rpm five minutes, then ultrasonic bath for twenty minutes. Peptide samples were centrifuged at 21,000 g for seven minutes at room temperature, quantified by microBCA assay (ThermoFisher Scientific), and then transferred to Waters Total Recovery autosampler vials for injection.

### Liquid Chromatography Mass Spectrometry and Data Analysis

Liquid chromatography mass spectrometry (LC-MS/MS) was performed in the Biomolecular Core (BioCORE) Facility at Western University Schulich School of Medicine and Dentistry, London, Canada. Peptide samples were injected onto a Waters Acquity UPLC M-Class system (Waters) coupled to a Q Exactive Plus hybrid quadrupole-Orbitrap mass spectrometer (ThermoFisher Scientific). Samples were trapped (5 μL min-1, 5 minutes) on a Symmetry C18 Trap Column, 5 μm, 180 μm × 20 mm (Waters) using 99% Mobile Phase A (H_2_O/0.1% formic acid (FA)), 1% Mobile Phase B (ACN/0.1% FA), then resolved on a Peptide BEH C18 Column, 130 Å, 1.7 μm, 75 μm × 250 mm (Waters) at 35°C and a flow rate of 300 nL min-1 using a nonlinear gradient (1-7% B, 1 minute; 7-23% B, 66.5 minutes; 23-35% B, 22.5 minutes; 35-98% B, 5 minutes). All liquid chromatography (LC) and mass spectrometry (MS) solvents used were of Optima LC/MS grade from Fisher Scientific.

The Q Exactive Plus mass spectrometer was controlled by Xcalibur software v4.0 (ThermoFisher Scientific). Data were acquired in data-dependent acquisition mode (DDA) using a FT/FT/HCD (Fourier Transform/higher-energy collision dissociation) Top 12 scheme. Survey scans (MS1) were acquired from m/z 375 to 1,500 at a resolution of 70,000 with AGC (automatic gain control) set to 3 e6 and a maximum injection time of 250 ms. Multiply charged peptide ions were isolated using a quadrupole isolation window of m/z 1.2 and were fragmented using HCD with a NCE (normalized collision energy) set to 25%. Fragment ion scans (MS2) were acquired at a resolution of 17,500 with AGC set to 2 e5 and a maximum injection time of 64 ms. Dynamic exclusion was set to 30 s, and lock mass ion was enabled at m/z 445.120025.

Raw data were processed using PEAKS 13 (Bioinformatics Solutions Inc.) and searched against all UniProtKB Homo Sapiens reviewed sequences (downloaded Sept. 12, 2025). Precursor and fragment ion mass tolerance was set to 10 ppm and 0.02 Da, respectively. Specific cleavage with trypsin was selected with a maximum of 3 missed cleavages, peptide length of 4-45 residues, and variable modifications of carbamidomethylation (C), deamidation (NQ), and oxidation (M), with a maximum of 5 variable modifications allowed per peptide. Peptide and protein group false discovery rates (FDR) were set to 1% for database searches. For label-free quantitation, peptide features were matched by automated retention time alignment with a mass error of 10 ppm and feature intensity ≥1 e5, and filtered for average area ≥1 e4, Quality score ≥5, and charge state of 1-6. Proteins were filtered to use ≥3 unique peptides with outlier and modified peptide exclusion, up to top 3 peptide for quantitation, a significance ≥15 (ANOVA), and fold-change ≥1.2 or ≤0.83 across n ≥ 5 biological replicates.

### A375-P Tumour Growth, Viability, and Metastasis in Xenograft Chick Chorioallantoic Membrane Assay (Chick-CAM)

Fertilized chicken eggs obtained from McKinley Hatchery (St. Mary’s, ON, Canada) were incubated in a 1502 Sportsman Rotating Incubator (GQF Manufacturing) maintained at 37°C for three days. The chick embryos were then transferred into a UV irradiated 100 mL weighing dish (Fisher Scientific, Cat# 08-732- 113) and covered with square disposable petri dish lids (Fisher Scientific, Cat# 0875711A) where they were further incubated in a 1550 Sportsman Hatching Incubator (GQF Manufacturing) for seven days at 37°C. SC and PANX1 KO A375-P cells were transduced with LV-tdTomato-Luc2 (tdT-Luc2) as described previously (77,78). Following 10 days of incubation, one million cells/chick were resuspended in 1x PBS, combined at a 1:1 ratio with phenol red-free Matrigel (Corning #356237) and implanted onto a branching vessel of the chorioallantoic membrane. tdT-Luc2 SC and PANX1 KO tumours were treated topically with 10 µM SPIR or vehicle control for seven days as described previously (22,30). Six days following implantation, tumours were imaged using an Eclipse Ni-E Fluorescence Microscope (Nikon) with a LU Plan Fluour EPI 5x objective to assess the visible surface area of tumours which was quantified using Fiji ImageJ Version 2.16.0/1.54p. On day seven-teen, endpoint images were captured using a digital camera and tumour viability was assessed with bioluminescent imaging (BLI) on an IVIS Lumina XRMS (PerkinElmer) following the topical application of D-luciferin (MediLumine, Cat# 222PS). Chicks were sacrificed and dissected for consideration of micro-metastasis to the liver, brain, and lungs as described previously (30). RNA was isolated using the RNeasy mini kit (Qiagen, Cat#74104) as per manufacturer’s directions, reverse transcribed using a High-Capacity cDNA Reverse Transcriptase Kit (Thermo Fisher Scientific, Cat #4368814) and cDNA was synthesized with a T100 Thermal Cycler (BioRad). RT-qPCR was performed in triplicate using a CFX Connect Real-Time PCR Detection System (Bio-Rad) on cDNA combined with SsoAdvanced Universal SYBR® Green Supermix (Bio-Rad, #1725274) and the following primer mix (human GAPDH Forward 5’-AATCCCATCACCATCTTCCA-3’ and Reverse 5’- TGGACTCCACGACGTACTCA-3’, or chicken GAPDH Forward 5’- GAGGAAAGGTCGCCTGGTGGATCG-3’ and Reverse 5’-GGTGAGGACAAGCAGTGAGGAACG-3’, along with D-Luciferase Forward 5’-GCTGGTGCCAACCCTATTCT-3’ and Reverse 5’- TCATCCCCCTCGGGTGTAAT-3’). In all cases, DNA from a tdT-Luc2 SC or PANX1 KO A375-P tumour was used as a positive control.

### Statistical Analysis

Data presented is depicted as mean ± the standard error of the mean (S.E.M.) when n>5 and mean when n<5. Data was analyzed using a Two-Tailed Unpaired T-Test, Two-Tailed Paired T-Test, Ordinary One-way ANOVA with Tukey post-test, or Two-way ANOVA with Tukey post-test as indicated in the figure captions. In all instances, experimental replicates are denoted as ‘n=’ and the number of biologically independent CRISPR clones assessed is denoted ‘N=’. Statistical analysis was performed using GraphPad Prism Version 11 (GraphPad Software, San Diego, California, https://www.graphpad.com).

### Data Availability Statement

The raw bulk RNA sequencing data generated in this study is publicly available in the Gene Expression Omnibus (GEO) repository at GSE348006. Proteomic data will be available through PRIDE database.

## RESULTS

### PANX1 is Ablated in A375-P Cells when Targeting Exon 1 of the *PANX1* Gene

To explore the mechanistic underpinnings governing the role of PANX1 in melanoma we employed CRISPR-Cas9 gene editing technology to characterize how the ablation of *PANX1* modulates melanoma cell properties. A375-P cells were nucleofected with three guide RNA (gRNA) directed to the first exon of the *PANX1* gene which encodes for the amino terminal region of the PANX1 protein **(Figure 1A-B)**. Following nucleofection, six scrambled (SC) and twelve PANX1 knockout (KO) A375-P clones were isolated, and the ablation of *PANX1* was validated via immunoblotting **(Figure 1C, E) (Supplementary Figure 1A-B)**. Quantification of immunoblots revealed a significant reduction in the levels of PANX1 when assessed by labeling with antibodies that target both the amino- and carboxy-termini of PANX1 **(Figure 1D, F)**. Moreover, we observed the complete loss of cell surface PANX1 staining in PANX1 KO A375-P cells confirming that PANX1 was successfully ablated **(Figure 1G) (Supplementary Figure 1C)**. Given the successful ablation of *PANX1*, we next sought to assess the consequences on melanoma cell growth over time. We found that the deletion of *PANX1* did not alter A375-P cell growth, although there was some variability between the twelve independently isolated PANX1 KO clones **(Supplementary Figure 1D, E)**.

**Figure 1:**
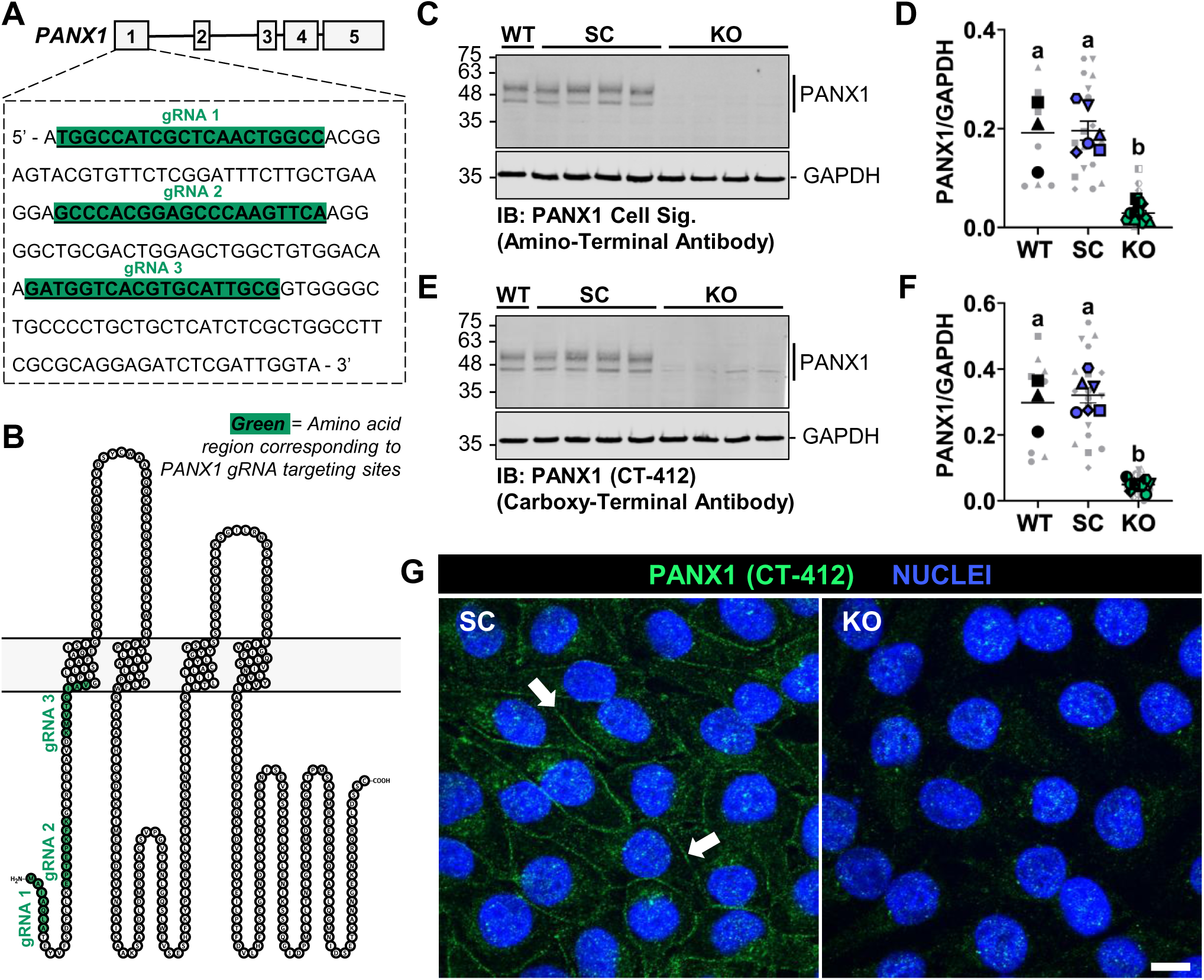
Successful CRISPR-Cas 9 mediated ablation of PANX1 in A375-P cells. **(A)** The first exon of the human *PANX1* gene (**B)** corresponding to the amino-terminal region of the PANX1 protein was targeted with three distinct guide RNA (gRNA) to ablate *PANX1* in A375-P cells using CRISPR-Cas9 gene editing technology. The ablation of *PANX1* was validated via immunoblots probing wild-type (WT) (n=3), scrambled control (SC) (N=6, n=3), and PANX1 knockout (KO) (N=12, n=3) A375-P cells with distinct antibodies directed against the **(C)** amino terminus (PANX1 Cell Sig.) or **(E)** carboxy-terminus (PANX1 (CT-412)) of the PANX1 protein. Quantification confirms that PANX1 KO A375-P cells exhibit a significant reduction in PANX1 protein levels when assessed with an **(D)** amino terminal (PANX1 Cell Sig.) or **(F)** carboxy-terminal (PANX1 (CT-412)) PANX1 antibody. For SC and PANX1 KO A375-P cells, blue or green data points denote the mean for each independent CRISPR clone while the grey data points depict each independent biological replicate. For WT A375-P cells, black data points denote the mean for each biological replicate while the grey data points depict technical replicates. Ordinary One- way ANOVA with Tukey post test; error bars are mean ± s.e.m. **(G)** PANX1 KO A375-P cells (N=4, n=3) exhibit a loss of cell surface PANX1 labelling (PANX1 (CT-412) antibody; green) when compared to SC (N=4, n=3) A375-P cells (arrows). Nuclei were stained with Hoechst 33342 (blue). Scale bar = 10 µm.

### Hallmark E2F Targets and G2/M Checkpoint Gene Sets are Negatively Correlated with the Targeted Deletion or Inhibition of PANX1

To comprehensively assess how the deletion of *PANX1* versus inhibition of its channel function impacts the transcriptome of melanoma cells, SC and PANX1 KO A375-P clones were treated with SPIR and subjected to bulk RNA sequencing **(Figure 2A)**. Principal component analysis (PCA) was used to validate clustering of samples according to each experimental condition **(Supplementary Figure 2A-C)**. In vehicle control conditions, we observed that the deletion of *PANX1* in A375-P cells resulted in 1,795 differentially expressed genes (DEGs) (968 downregulated and 827 upregulated) compared to SC cells when using a threshold significance of 0.05 and a Log2(Fold Change) of |1.5| **(Figure 2B, C)**. Comparatively, when PANX1 channels were inhibited with SPIR in SC cells, we identified 799 DEGs (405 downregulated and 394 upregulated) compared to vehicle treated SC cells **(Figure 2D, E).** Whereas the same treatment in PANX1 KO A375-P cells resulted in 968 DEGs (611 downregulated and 357 upregulated) compared to vehicle treated KO cells **(Figure 2F, G)**. The ablation and inhibition of PANX1 resulted in distinct changes to the transcriptional profile of A375-P cells with only 263 of the same genes being differentially expressed in both experimental paradigms (116 downregulated and 147 upregulated) **(Figure 2H-J)**. However, gene set enrichment analysis (GSEA) revealed that cell cycle-associated gene sets including Hallmark E2F Targets and G2/M Checkpoint were positively correlated in vehicle treated SC cells when compared to both the ablation and inhibition of PANX1 with SPIR **(Figure 2K, L) (Supplementary Figure 3)**. Surprisingly, despite lacking PANX1 channels to inhibit, the same trend was also observed in vehicle treated PANX1 KO cells when compared to PANX1 KO cells treated with SPIR **(Figure 2M) (Supplementary Figure 3)**. The transcript expression of Hallmark E2F Targets and G2/M Checkpoint genes were similarly downregulated following the deletion and inhibition of PANX1 channel function **(Figure 2N, O)**. Although, the greatest difference was observed when PANX1 KO A375-P cells were treated with SPIR, suggesting that the loss of PANX1 may sensitize A375-P cells to the pleotropic effects of SPIR which further exacerbates the dysregulation of Hallmark E2F Targets and G2/M Checkpoint genes.

**Figure 2:**
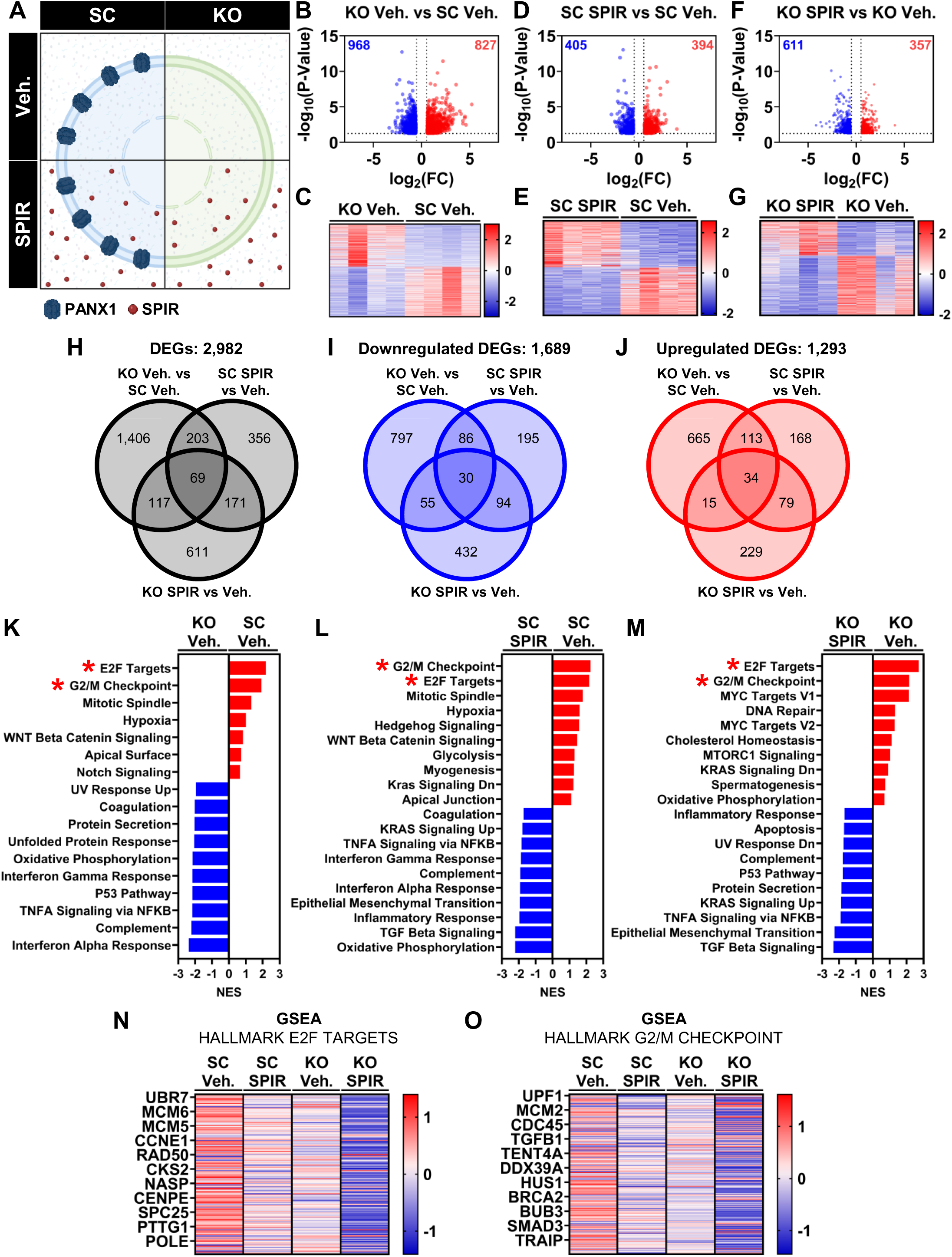
Cell cycle pathways are negatively enriched with the ablation and inhibition of PANX1. **(A)** Depiction of experimental conditions examined. Scrambled control (SC) and PANX1 KO A375-P cells were treated with PANX1 inhibitor spironolactone (SPIR; 10 μM or vehicle control) for 96 hours and subsequently isolated for bulk RNA sequencing. **(B, D, F)** Volcano plots and **(C, E, G)** heatmaps depicting the differentially expressed genes (DEGs) indicate differences the transcriptional profile of A375-P cells when PANX1 was deleted or inhibited with SPIR(N=4) (significance of 0.05 and a Log2(Fold Change) of |1.5|). **(H)** Across all experimental conditions, there were 2,982 differentially expressed genes (DEGs) with **(I)** 1,689 downregulated and **(J)** 1,293 upregulated. **(K-M)** Gene set enrichment analysis (GSEA) revealed that the deletion versus inhibition of PANX1 differentially affects various biological pathways, although cell cycle related pathways (red asterisk) were commonly disrupted (positively correlated gene sets (red) and negatively correlated gene sets (blue)). Specifically, the Hallmark E2F target, and G2/M checkpoint genes were positively correlated in vehicle treated SC cells when compared to the deletion of *PANX1* or treatment with SPIR. **(N, O)** Heatmap of all E2F target and G2/M checkpoint genes reveals that cell cycle related transcripts are downregulated following the deletion of *PANX1* and inhibition of PANX1 in SC A375-P cells, although the expression of these genes was further reduced when PANX1 deficient A375-P cells were treated with SPIR.

### PANX1 Inhibitors Reduce the Growth of Adherent SC and PANX1 KO A375-P Cells

To investigate the phenotypic consequences to cell growth, we challenged SC A375-P clones with the PANX1 inhibitor SPIR and found that the inhibition of PANX1 reduced cell growth over time, a finding that is consistent with previous studies **(Figure 3A-C)** (22,23,55). Similarly, we observed a significant reduction to cell growth over time when PANX1 KO A375-P cells were treated with SPIR which is expected based on our bulk RNA sequencing results where the most pronounced transcriptomic change occurred with the concurrent ablation of *PANX1* and treatment with SPIR. To assess whether this reduction in growth was due to differences in cell death, we assessed molecular indicators of endoplasmic reticulum (ER)stress (CHOP, BiP), autophagic flux (LC3-II/I), and apoptosis (cCASP3), and observed no statistical differences **(Supplementary Figure 4)**. Next, we monitored cell cycle progression using flow cytometry to assess whether the deletion of *PANX1* or treatment with SPIR disrupts cell proliferation. To mitigate the variability between distinct clonal populations, we proceeded to reduce our characterization to four independent SC (SC1, SC2, SC3, SC4) and PANX1 KO (KO1, KO2, KO3, KO4) A375-P clones which were pooled at an equal cell number to form a mixed clonal population. Selection of A375-P clones was determined based on immunoblots quantifying the abundance of PANX1, where selected SC clones exhibit similar levels to WT A375-P cells and PANX1 KO clones display the lowest levels of PANX1 signal **(Supplementary Figure 1A-C)**. Interestingly, we found that the ablation of *PANX1* shifted cell cycle distribution from G0/G1 towards G2/M **(Figure 3D)**. Conversely, treatment with SPIR resulted in an increased proportion of cells in G0/G1 with a reduced proportion of cells detected in S and G2/M, suggesting SPIR stalls A375-P cells at the G1/S checkpoint. To determine whether this is a unique effect of SPIR, we additionally treated SC and PANX1 KO A375-P cells with the PANX1 channel inhibitor PBN **(Figure 3E)**. Similarly to treatment with SPIR, PBN reduced the growth of SC and PANX1 KO A375-P cells without any indication of elevated ER stress, autophagic flux, or apoptosis **(Figure 3F, G) (Supplementary Figure 4)**. Analogous to the deletion of *PANX1*, PBN treatment increased the proportion of SC A375-P cells in G2/M with no differences detected at any cell cycle checkpoint when PANX1 KO cells were treated with PBN indicating that SPIR and PBN function through different cellular mechanisms **(Figure 3H)**. Cumulatively these findings indicate that PANX1 channel activity is likely involved in modulating cell cycle progression since the deletion of *PANX1* and inhibition with PBN analogously stall cell cycle progression at G2/M. Whereas, spironolactone stalls A375-P cells at the G1/S checkpoint in a PANX1 independent manner. Importantly, treatment with either SPIR or PBN did not change the levels or visually alter the cell surface localization of PANX1 in SC A375-P cells **(Supplementary Figure 5)**.

**Figure 3:**
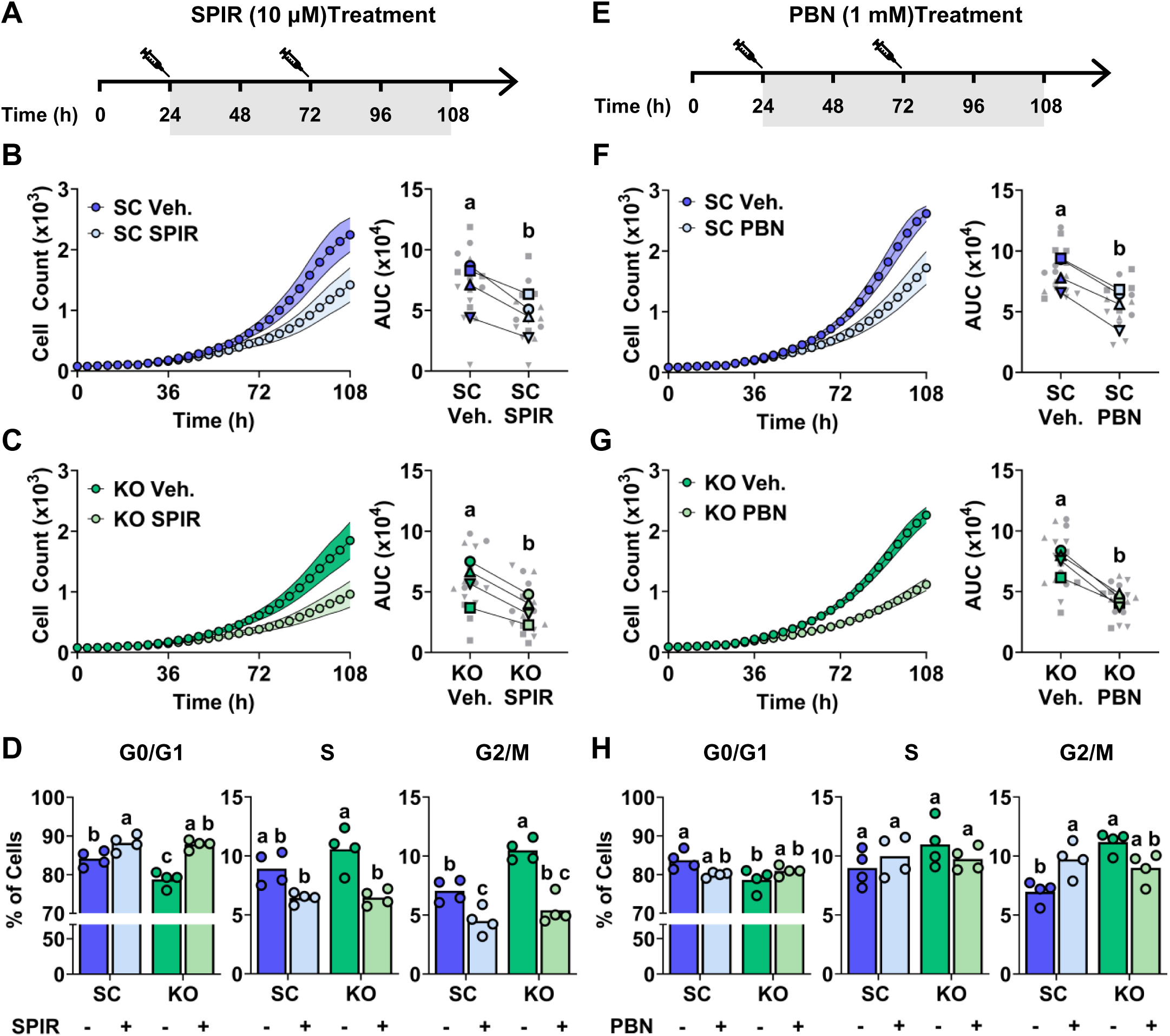
PANX1 inhibitors spironolactone and probenecid reduce the growth of adherent A375-P cells. Schematic illustrating the experimental timeline of PANX1 inhibition with **(A)** 10 μM SPIR or **(E)** 1 mM probenecid (PBN). SC (N=4, n=3) cell growth was assessed over 108 hours to examine how the inhibition of PANX1 channel function with **(B)** SPIR or **(F)** PBN impacts A375-P cell growth over time. Area under the curve (AUC) quantification of cell growth curves reveled that treatment with PANX1 channel inhibitors significantly reduced SC A375-P cell growth. Cell growth was significantly reduced when PANX1 KO A375-P cells were treated with either **(C)** SPIR or **(G)** PBN. Blue and green data points denote the mean AUC for each independent CRISPR clone, while the grey data points depict each independent biological replicate. Two-Tailed Paired T-Test. **(D;** SPIR, **H**; PBN) Flow cytometry revealed that the deletion of *PANX1* shifts cell cycle distribution from G0/G1 to G2/M in A375-P cells. Inhibition of PANX1 with PBN stalled SC A375-P cells at the G2/M checkpoint whereas SPIR treatment stalled both SC and PANX1 KO A375-P cells at the G1/S checkpoint, indicating that these PANX1 channel inhibitors likely reduce cell growth via distinct cellular mechanisms (n=4). Two-way ANOVA with Tukey post-test.

### PANX1 Promotes Spheroid Compaction and Adhesivity in Melanoma Cells

To assess the role of PANX1 in a melanoma model that better recapitulates the 3D architecture of tumours, we generated SC and PANX1 KO A375-P spheroid cultures. We assessed spheroid formation over 6 days and observed that SC A375-P cells spontaneously form compact, well-defined spheroids **(Figure 4A)**. The deletion of *PANX1* disrupts spheroid compaction, with PANX1 KO A375-P spheroids exhibiting poorly defined peripheral borders, numerous projections, and a significantly reduced circularity **(Figure 4B, C)**. Similarly, when SC A375-P spheroids were challenged with SPIR there was a significant reduction in circularity with spheroids morphologically, resembling the PANX1 KO condition **(Figure 4B, C)**. This finding suggests PANX1 channel function may be responsible for the regulation of spheroid compaction over time. When PANX1 KO A375-P spheroids were challenged with SPIR there was an exacerbated reduction in spheroid circularity and increase in overall spheroid area, indicating that the deletion of *PANX1* may sensitize A375-P cells to SPIR treatment which further destabilizes spheroid assembly **(Figure 4B-D)**.

**Figure 4:**
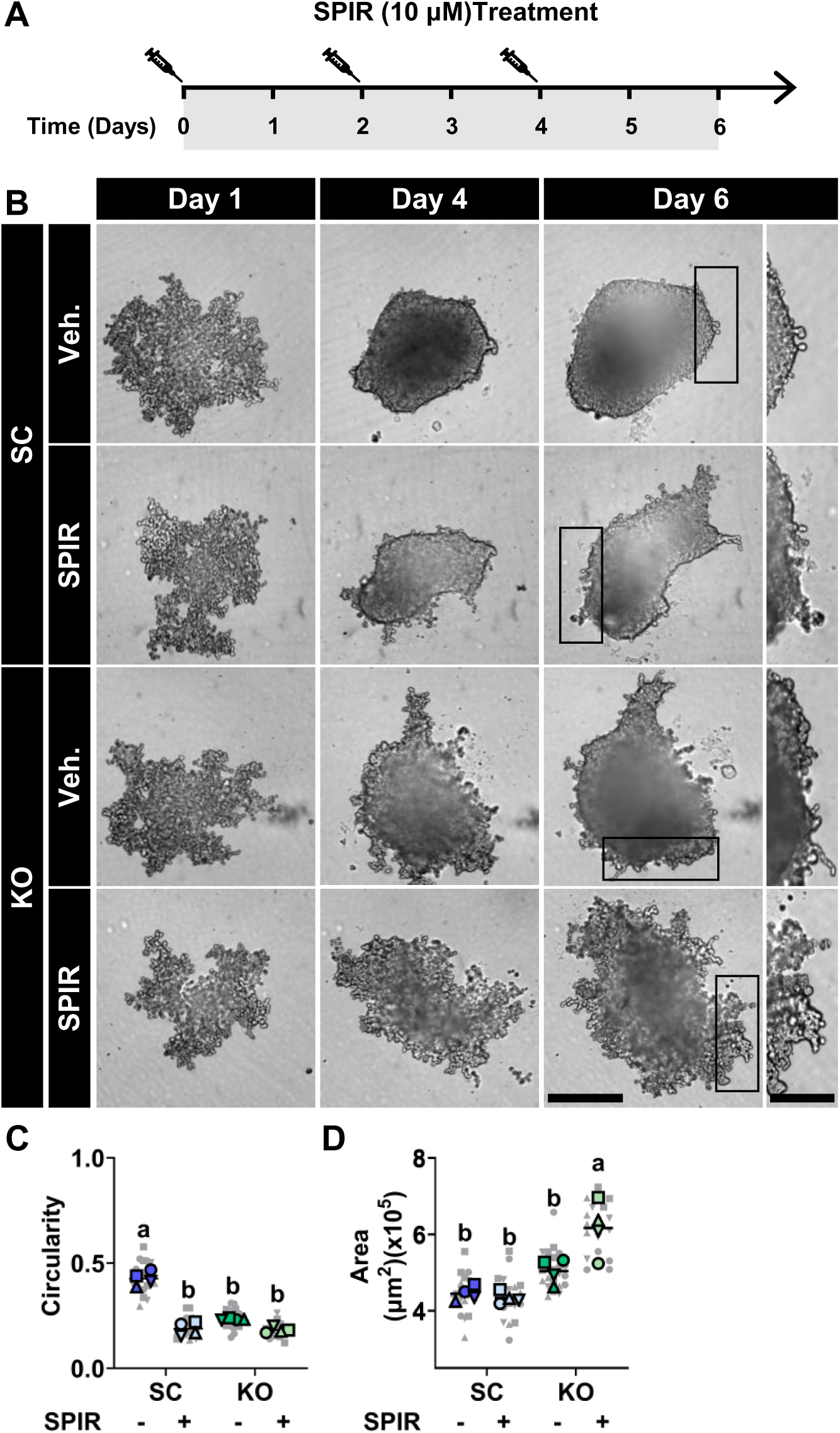
PANX1 targeting strategies disrupt spheroid morphology. **(A)** Schematic illustrating the spheroid experimental timeline of PANX1 inhibition with 10 μM SPIR. **(B)** Representative phase contrast images of SC and PANX1 KO A375-P spheroids (n=4) treated with 10 μM SPIR over a period of 6 days. Scale bar = 400 µm. The genetic ablation of *PANX1* and/or treatment with SPIR significantly disrupted spheroid compaction over time with spheroids exhibiting irregular borders relative to control conditions (magnified area; scale bar = 200 µm). **(C)** Quantification of spheroid circularity at endpoint revealed that deletion and/or inhibition of PANX1 reduced spheroid circularity in all experimental paradigms. **(D)** Spheroid *en face* area was quantified at endpoint, revealing PANX1 KO A375-P spheroid area significantly increased when challenged with SPIR. Blue and green data points denote the mean for each biological replicate while the grey data points depict technical replicates (4-6 spheroids per experimental condition). Two-way ANOVA with Tukey post-test.

Next, we sought to assess whether this change to spheroid compaction and circularity was due to differences in spheroid viability. Using calcein-AM, a cell permeable dye that labels metabolically active cells, we found no differences to spheroid viability following the deletion or inhibition of PANX1 **(Figure 5A, B)**. However, the ablation of *PANX1* sensitized spheroids to fragmentation following mechanical stress from pipetting, suggesting that the deletion of *PANX1* may compromise cell-cell adhesions within the spheroid **(Figure 5C)**. Moreover, Kyoto Encyclopedia of Genes and Genomes (KEGG) pathway analysis of bulk RNA sequencing revealed significant pathway enrichment of cell adhesion molecule genes with the genetic ablation of *PANX1*, as evidenced by an upregulation of cell adhesion molecule mRNA expression **(Figure 5D, E)** (67–69). To understand how PANX1 contributes to the structural integrity of A375-P spheroids we immunolabelled spheroids to assess the localization and distribution of PANX1 signal. We observed PANX1 signal uniformly distributed throughout the core and periphery of SC A375-P spheroids when assessed using two distinct PANX1 antibodies **(Figure 5F, G)**. Altogether this suggests that PANX1 may act as a scaffold protein to stabilize cell-cell contacts within spheroids and ultimately support the structural integrity of A375-P spheroids.

**Figure 5:**
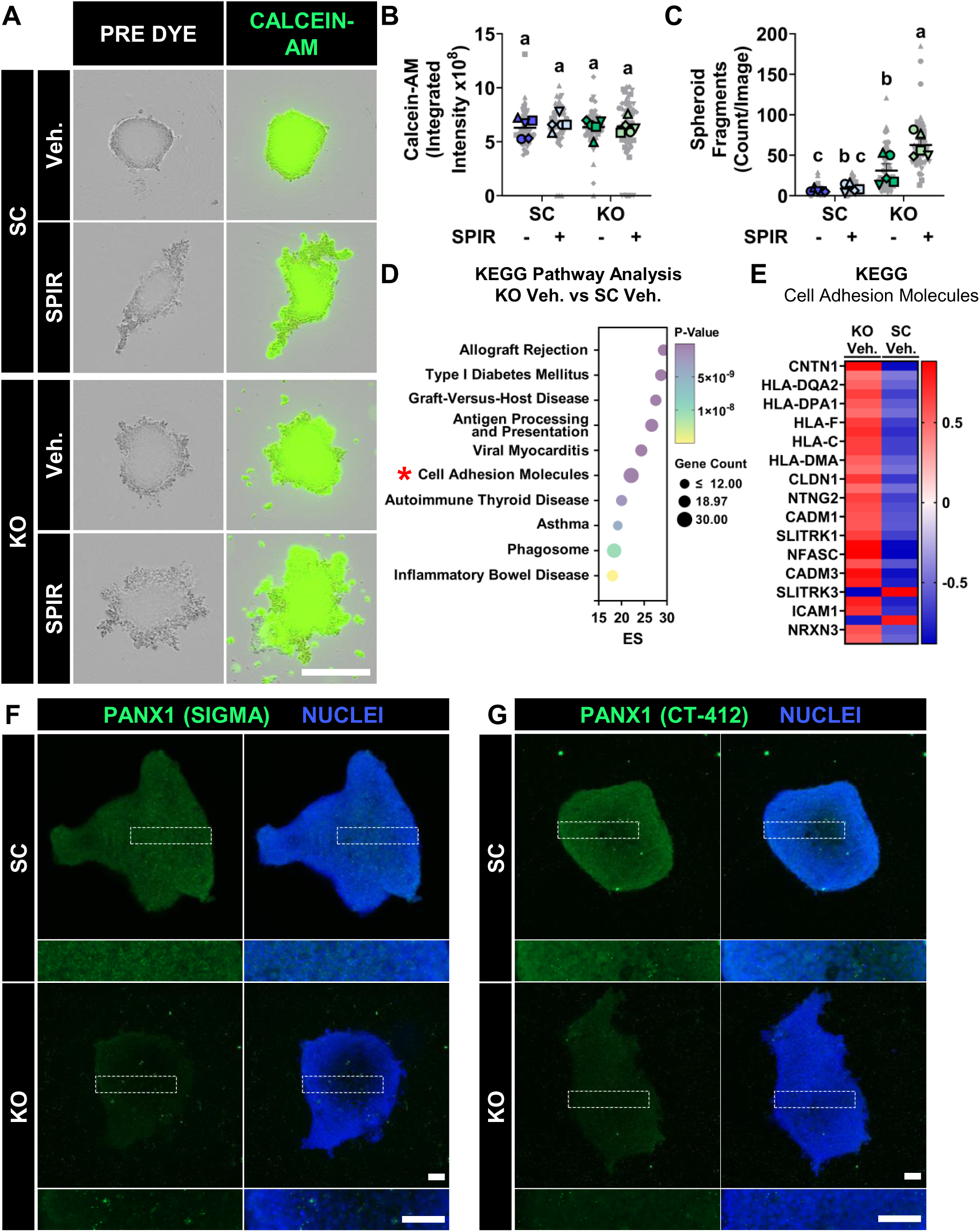
Melanoma spheroid adhesion is compromised following the ablation of *PANX1*. **(A)** Representative phase contrast images of SC (n=5) and PANX1 KO A375-P spheroids (n=5) before (pre-dye) and after labelling with calcein-AM (green) dye to assess spheroid viability. Scale bar = 800 µm. **(B)** The deletion or inhibition of PANX1 does not alter spheroid viability as indicated by calcein-AM fluorescence intensity (green; labels metabolically active cells). **(C)** The genetic ablation of *PANX1* significantly increased spheroid fragmentation following mechanical stress. Blue and green data points denote the mean for each biological replicate while the grey data points depict technical replicates (8-12 spheroids per experimental condition). Two-way ANOVA with Tukey post-test. **(D)** Kyoto Encyclopedia of Genes and Genomes (KEGG) pathway analysis of bulk RNA sequencing (N=4) from adherent SC and PANX1 KO A375-P cells revealed pathway enrichment of cell adhesion molecules with the genetic ablation of PANX1. **(E)** Heatmap of differentially expressed genes within the KEGG Cell Adhesion Molecule Pathway reveals that cell adhesion molecule transcripts are upregulated following the deletion of *PANX1*. **(F, G)** PANX1 signal (green) is uniformly distributed throughout the core and periphery (magnified area) of SC A375-P spheroids (n=3) where it is detectable within intracellular compartments and at the surface of individual cells when assessed with two distinct carboxy-terminal targeted PANX1 antibodies. Nuclei were stained with Hoechst 33342 (blue). Scale bar = 100 µm.

### Multiple Cell Adhesion Proteins Co-Immunoprecipitate with PANX1

Mechanistically, we next sought to assess whether the phenotypic changes observed with PANX1 KO A375-P spheroids could be explained due to the disruption of key stabilizing interactions resulting from PANX1 ablation. As such, PANX1 was immunoprecipitated from SC A375-P cells using a carboxy-terminal PANX1 antibody which isolates PANX1 in its different glycosylation states appearing as multiple bands between 45-60 kDa. In contrast, PANX1 was not detected when immunoprecipitated from PANX1 KO A375-P cells, with only the heavy chain immunoglobulin G (IgG) being detected at 50-55 kDa which is similar to the isotype IgG control immunoprecipitation reactions **(Figure 6A)**. After confirming the successful isolation of PANX1, subsequent immunoprecipitations were performed and co-immunoprecipitated proteins were identified using liquid chromatography mass spectrometry. Proteomic analysis revealed enrichment of 76 proteins, including PANX1 in SC A375-P cells compared to PANX1 KO cells **(Figure 6B)**. We cross referenced the 75 potential PANX1 interacting partners with the publicly available BioGRID database and found that only 7 of the proteins we identified (GNAS, HSD17B12, TMCO1, ATP1A1, ACSL3, COPA, ARF1) are currently recorded as PANX1 interacting partners (79). Importantly, PANX1 was identified as the most significantly enriched protein in our screen validating the successful immunoprecipitation of PANX1 across all 5 of our technical replicates **(Figure 6B)**. We performed gene ontology (GO) pathway analysis on the 75 putative PANX1 interacting partners which revealed an enrichment in proteins that are involved in cell adhesion molecule binding, cytoskeletal regulation, and intracellular transport **(Figure 6C)**. Protein-containing complex binding was identified as the most enriched molecular function for the identified proteins which includes β-actin, a well-established PANX1 interacting partner **(Figure 6D)**(36,80). Taken together, this enrichment suggests that PANX1- mediated interactions may contribute to cellular organization and tumour stability in 3-dimensional tumor spheroids.

**Figure 6:**
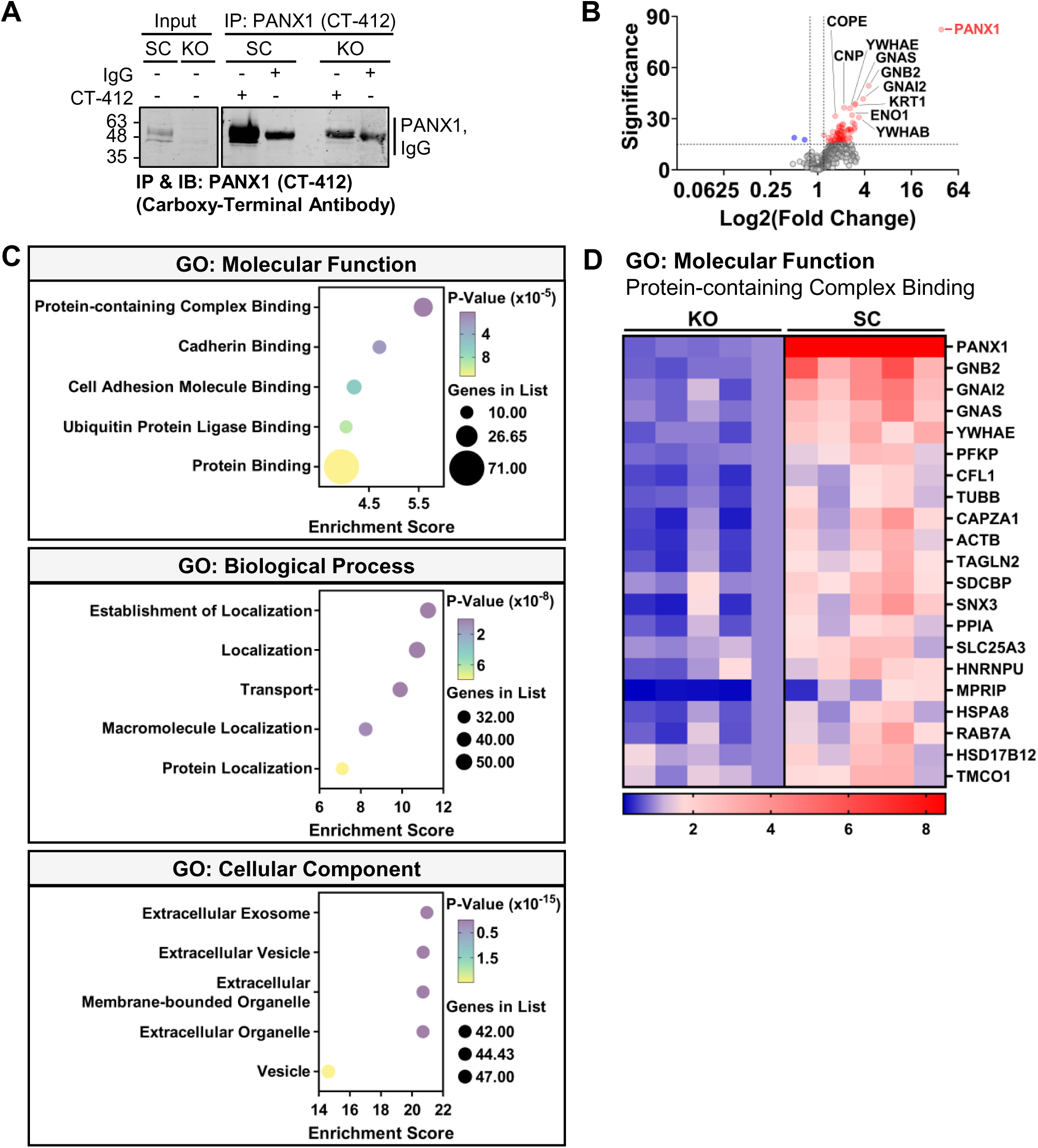
Mass spectrometry analysis revealed enrichment in cell adhesion and trafficking proteins co-immunoprecipitating with PANX1. **(A)** PANX1 was immunoprecipitated using the PANX1 (CT-412) antibody from SC A375-P cells and PANX1 KO cells as a control. To confirm the successful isolation of PANX1, an additional immunoprecipitation was run in parallel and visualized via immunoblotting. Rabbit immunoglobulin G (IgG) was used as negative control (n=1). (B) Liquid chromatography mass spectrometry identified significant enrichment of 75 proteins in SC A375-P cells that were co-immunoprecipitated with PANX1 (n=5). **(C)** Gene ontology (GO) pathway analysis of the 75 putative PANX1 interacting partners revealed enrichment in proteins involved in protein-protein binding and cellular trafficking. **(D)** Heatmap of the 21 immunoprecipitated proteins that belong to the Protein- containing Complex Binding node which was identified as the most enriched functional node to describe the molecular activities of the proteins that co-immunoprecipitate with PANX1.

### The Deletion of *PANX1* Reduces Tumour Viability and Disrupts the Spatial Organization of Tumours in a Xenograft Chick Chorioallantoic Model

As the deletion of *PANX1* sensitized SPIR treated PANX1 KO A375-P cells to dissociate when challenged with mechanical stress, we questioned whether this would facilitate increased migration. Using a scratch wound closure assay, we found no changes to cell migration upon the inhibition of PANX1 with channel inhibitors SPIR or PBN when compared to vehicle-treated SC cells **(Supplementary Figure 6)**. However, as migration was assessed in two dimensions this may not recapitulate the complexity of a three-dimensional system. As such, we assessed spontaneous metastasis in a xenograft chick chorioallantoic model to evaluate the migratory capacity and viability of A375-P cells when extracellular matrix interactions are supported with Matrigel. First, SC and PANX1 KO A375-P cells were transduced with LV-tdTomato-Luc2 (tdT-Luc2) to express tdTomato and firefly luciferase as reporters for downstream analysis. tdT-Luc2 SC and PANX1 KO A375-P cells were grafted onto a branching vessel and subsequently treated topically with 10 µM SPIR daily to assess how PANX1 targeting strategies impact melanoma tumour growth and metastasis *ex vivo* **(Figure 7A)**. To investigate whether the deletion of *PANX1* enhanced migration and invasion, we assessed spontaneous metastasis to the embryonic brain, liver, and lungs and observed no transcript amplification in any of the assessed organs regardless of treatment condition (data not shown). However, bioluminescence imaging (BLI) of the primary tumour at endpoint revealed a genotype effect (p=0.05) with a trend towards reduced overall viability in tumours due to the loss of PANX1 **(Figure 7B-C)**. At endpoint, tumours were excised and weighed to evaluate overall tumour growth, but no differences in tumour weight were observed when PANX1 was deleted or inhibited with SPIR **(Figure 7D)**. However, *ex vivo* fluorescence imaging revealed a genotype effect (p=0.05) where the ablation of *PANX1* significantly reduced the visible *en face* tumour area relative to controls **(Figure 7E-F)**. Thus, suggesting that the ablation of *PANX1* reduces lateral tumour growth which may reflect a difference in the spatial organization of A375-P tumours.

**Figure 7:**
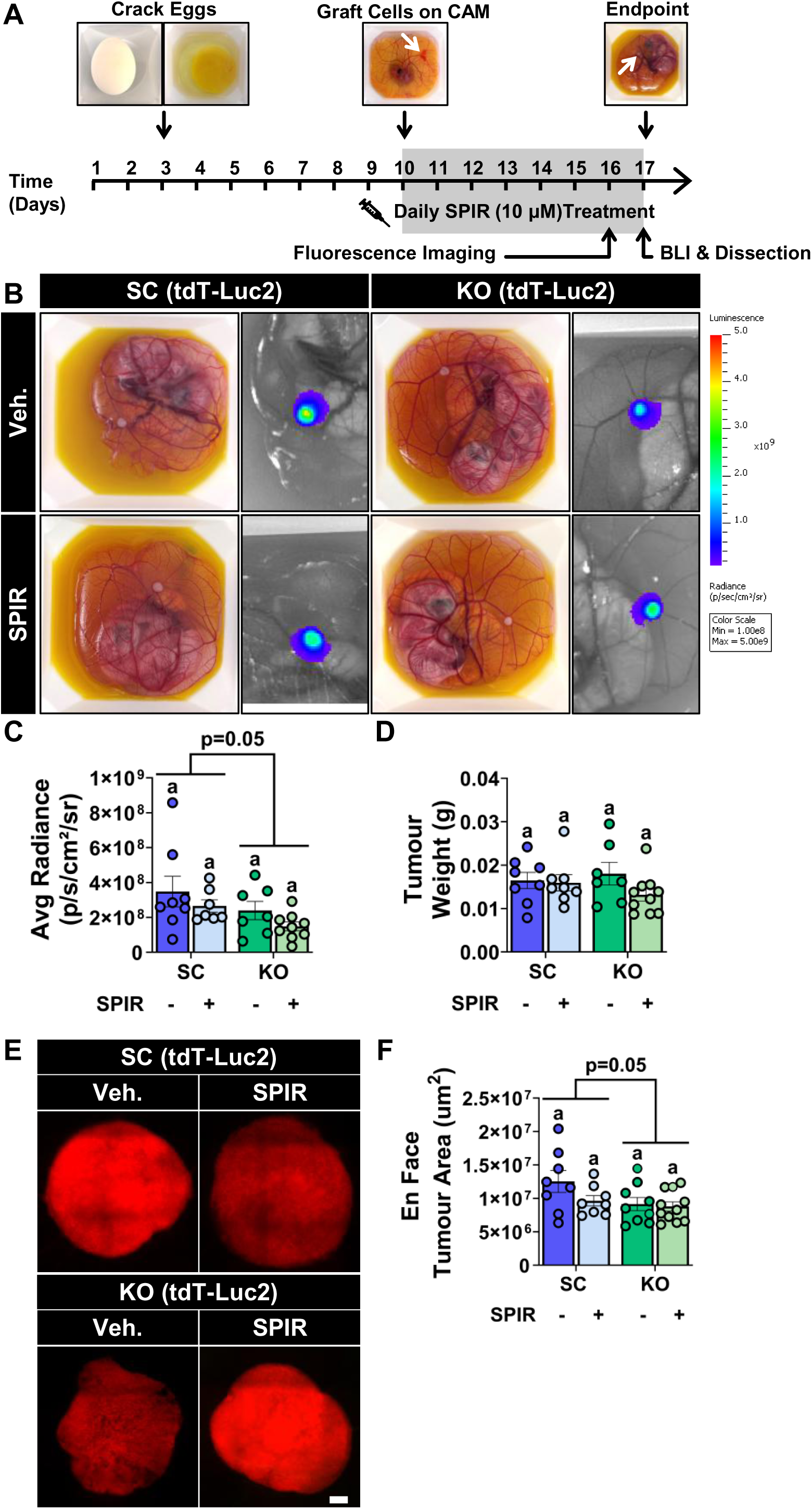
The deletion of PANX1 reduced viability and *en face* area of A375-P tumours *ex vivo.* **(A)** Schematic depicting the experimental timeline for the A375-P xenograft chick chorioallantoic model (CAM). Following incubation, SC and PANX1 KO A375-P tumours were treated with 10 μM SPIR or vehicle control daily for seven days. Prior to implantation, SC and PANX1 KO A375-P cells were transduced with LV-tdTomato-Luc2 to express tdTomato and firefly luciferase to enable visualization with fluorescence and bioluminescence imaging (BLI), respectively, before endpoint. **(B)** Representative photograph with corresponding bioluminescence radiance levels for SC and PANX1 KO tumours treated with SPIR or vehicle control at endpoint. Scale bar depicts radiance intensity (p/sec/cm^2^/sr). **(C)** Quantification of peak radiance at endpoint reveals a genotype effect where there is a reduction in overall tumour viability due to the loss of PANX1 although there are no statistical differences in viability between each experimental group (n=7-9). **(D)** At endpoint there was no measurable change to overall tumour weight following the deletion or pharmacological inhibition of PANX1 (n=7-10). **(E)** Representative *en face* fluorescence images of SC and PANX1 KO tumours treated with SPIR or vehicle control. Scale bar = 500 µm. **(F)** Quantification of *en face* tumour area revealed a genotype effect where the deletion of PANX1 reduced the visible area of tumours *ex vivo* (n=8-11). Two-way ANOVA with Tukey post-test. Error bars are mean ± s.e.m.

## DISCUSSION

Metastatic melanoma remains a deadly malignancy despite significant advancements in the development of targeted therapies and immunotherapies (7–9). Our previous findings provided some insight regarding the mechanisms by which PANX1 contributes to melanoma pathogenicity, since we have shown that abolishing PANX1 channel function via pharmacological inhibition or silencing transcript expression with shRNA significantly reduces melanoma cell properties such as growth, migration, and invasion *in vitro* and *ex vivo* (22,30). This work raises questions as to how channel-dependent and - independent roles of PANX1 function in parallel to drive cellular properties associated with melanoma progression. Here we demonstrate that the ablation of *PANX1* shifted cell cycle distribution towards G2/M, which is consistent with our transcriptomic data showing the downregulation of E2F target and G2/M checkpoint genes following the deletion or inhibition of PANX1 channel function using spironolactone. Despite this, the deletion of *PANX1* did not alter A375-P cell growth *in vitro* nor elevated molecular indicators of ER stress, autophagic flux, or apoptosis, suggesting that the loss of PANX1 may impair mitotic entry potentially due to DNA damage or mitotic spindle disruption. Conversely, treatment with spironolactone reduced melanoma cell growth and stalled cell cycle progression at the G1/S checkpoint regardless of PANX1 protein expression. Moreover, cell cycle transcriptomic changes were exacerbated when PANX1 deficient cells were challenged with spironolactone highlighting a previously undefined synergy between PANX1 loss and spironolactone. We revealed that the deletion and inhibition of PANX1 compromises spheroid compaction, although this phenotype was worsened when PANX1 KO cells were challenged with spironolactone further demonstrating a synergistic relationship between these targeting strategies which could be exploited as an effective strategy to induce synthetic lethality in melanoma. PANX1 was found to stabilize spheroid cell-cell contacts which may be driven in part by its interactions with various components of the actin cytoskeleton. When assessed *ex vivo* the loss of PANX1 reduced *en face* tumour area and viability highlighting a therapeutic benefit of targeting PANX1.

Our transcriptomic analysis revealed that E2F target and G2/M checkpoint genes are downregulated with the targeted deletion of *PANX1* and inhibition of PANX1 channel function. Genes in the E2F target pathway are critical for DNA replication and cell cycle progression whereas G2/M checkpoint genes regulate cell division in response to DNA damage (81–83). We found that the CRISPR- Cas9 mediated ablation of *PANX1* shifted cell cycle distribution from G0/G1 towards G2/M which may be indicative of cellular stress or delayed mitotic entry as adherent cell growth was unchanged. Consistent with these findings, when control A375-P cells were treated with probenecid the proportion of cells occupying the G2/M phase significantly increased unlike when PANX1 KO cells were challenged with probenecid which resulted in no changes to cell cycle distribution. Taken together, our findings suggest PANX1 channel activity is involved in modulating cell cycle progression as the deletion of *PANX1* and inhibition with probenecid exhibited similar trends. However, despite no changes to cell cycle distribution when PANX1 KO cells were treated with probenecid, we observed a significant reduction in cell growth indicating that although the proportion of the cells in each phase remains consistent the overall duration of cell cycle progression in PANX1 KO cells is reduced with probenecid treatment. In contrast, we found that treatment with the PANX1 inhibitor spironolactone stalled A375-P cells at the G1/S checkpoint regardless of PANX1 expression. Thus, the growth deficits observed with spironolactone may not be entirely driven by the inhibition of PANX1 channel function, a finding that is unsurprising given spironolactone is a repurposed drug. Moreover, evading G2/M arrest is being investigated as a mechanism to achieve synthetic lethality in melanoma by inducing genomic instability (84,85). Thus, in addition to its PANX1 channel inhibitory function spironolactone may be a potential synthetic lethal agent through its pleotropic effects. Although originally developed as a steroidal mineralocorticoid receptor antagonist, spironolactone is now well regarded as a multi-target drug. For example, spironolactone was found to antagonize androgen and progesterone receptors at differing affinities (86,87). Moreover, the anti-adrenergic effect of spironolactone has been well accepted clinically leading to its off-label use for the treatment of dermatological conditions such as acne and hirsutism (88,89). However, the off-target effects of spironolactone are not limited to nuclear receptor family members, since cryogenic electron microscopy studies have revealed that spironolactone forms hydrophobic interactions within the central cavity of the human organic cation transporter 1 (PDB ID: 8JU0 & 8JTZ) leading to functional inhibition of the channel (90). Additionally, in a small molecule screen, spironolactone was identified as a nucleotide excision repair inhibitor that induces the proteasomal degradation of the DNA helicase xeroderma pigmentosum group B (XPB; encoded by ERCC3) (91–93). XPB is a core component of the transcription factor II H complex (TFIIH) and is essential for transcription initiation and DNA repair mechanisms (94,95). Thus, it is possible that the cell cycle changes we observed when PANX1 KO cells were challenged with spironolactone could be explained by the pleotropic effect of spironolactone on XPB preventing cells from undergoing transcription initiation and DNA repair mechanisms. Interestingly, transcriptomic analysis of patient skin cutaneous melanoma biopsies from The Cancer Genome Atlas indicates a moderate statistically significant positive correlation (Pearson: p−value = 7.5e−14, R = 0.34) between *PANX1* and *ERCC3* transcripts, highlighting a potential functional interplay between these distinct genes which could be investigated in future studies **(Supplementary Figure 7)** (96).

While adherent 2-dimensional monolayer cultures can be informative, they fail to recapitulate the complex network of cell-cell and cell-matrix interactions within a 3D tumour where cancer cell interactions with other cancer cells and the microenvironment is more complex. Notably, we found that PANX1- deficient cells exhibited reduced spheroid circularity with multiple protrusions projecting from the spheroid. This morphological change was phenocopied when cells were treated with spironolactone to inhibit PANX1 channels, implicating PANX1 channel function in the modulation of melanoma spheroid integrity. Similar to our findings, in patient-derived high-risk neuroblastoma cells, the inhibition of PANX1 with probenecid or carbenoxolone inhibits spheroid growth over time (23). Similarly, in C6 rat glioma cells, the overexpression of PANX1 accelerates the compaction of tumor aggregates, an effect which was reversed with the addition of the PANX1 channel inhibitor carbenoxolone, thus highlighting a role for PANX1 in the growth and structural assembly of spheroids (97). While we demonstrate that PANX1 plays a role in supporting the structural integrity of melanoma spheroids, the most prominent morphological change was observed when PANX1 KO spheroids were treated with spironolactone. Unlike control conditions, PANX1 KO A375-P spheroids challenged with spironolactone assembled into loose aggregates with irregular borders, increased surface area, and an increased propensity to dissociate. Spheroid formation can be separated into three distinct phases, an initial loose aggregation driven by integrin binding to the extracellular matrix (ECM), followed by cadherin-mediated spheroid compaction, and finally outwards growth where spheroids establish oxygen and nutrient gradients (98–100). It is possible that the ablation of *PANX1* compromises the efficacy of spheroid compaction, and this vulnerability can be exacerbated by the pleiotropic effects of spironolactone which corroborates other findings involving spironolactone treatment in cancer. For example, spironolactone treatment was shown to impair cancer cell survival when administered with Phleomycin in U2OS cells (101). Consistent with these findings, in a bladder carcinoma xenograft model, the combination of spironolactone with either cisplatin or carboplatin significantly impaired tumour growth as well as enhanced the treatment response following chemoresistance (102). Similarly, the combinatorial treatment of spironolactone with melphalan significantly reduced the viability of patient-derived multiple myeloma cell lines when compared to either treatment alone, suggesting spironolactone sensitizes cells to alkylating agents, an effect that was lost when assessed in F99S/R425X ERCC3 mutant cell lines (103). Interestingly, a Phase 1b/2a clinical trial (NCT Number: NCT07431216) is actively seeking to investigate the combinatorial treatment of spironolactone with the small molecule alkylating agent STAR-001(LP-184), in patients with IDH wild-type glioblastoma or IDH mutant grade 4 astrocytoma following success in preclinical xenografts (104). Our results suggest that spironolactone treatment could similarly be used to exploit the intrinsic vulnerabilities caused by the ablation of *PANX1* in melanoma to induce a synthetic lethal effect, although further research is warranted to understand the interplay between PANX1 and spironolactone along with its principal metabolites (49).

Given all three of our experimental conditions altered spheroid morphology, we questioned whether this could reflect a change to spheroid viability. In keeping with our adherent cell model, no changes to spheroid viability following the ablation or pharmacological inhibition of PANX1 were observed. We found however, that the deletion of *PANX1* destabilized cell-cell contacts within melanoma spheroids resulting in enhanced fragmentation following mechanical stress. No differences to spheroid fragmentation were observed upon the inhibition of PANX1 channels with spironolactone leading us to speculate that PANX1 stabilizes the integrity of spheroid cell-cell contacts via channel-independent means. Mechanistically, KEGG pathway analysis of our bulk RNA sequencing data revealed significant enrichment in cell adhesion molecule genes, implicating PANX1 as a potential player in maintaining the structural integrity of cell-cell contacts in melanoma. In keeping with this finding, we investigated PANX1 interacting partners and found that PANX1 predominantly co-immunoprecipitated proteins involved in cell adhesion molecule binding, cytoskeletal regulation, and intracellular transport. As expected, we identified β-actin (ACTB) a well-established PANX1 interacting partner and F-actin-capping protein subunit alpha-1 (CAPZA1) as co-immunoprecipitated proteins in our screen (36,80,105). It has been suggested that the interaction between PANX1 and actin related protein 3 (ARP3) modulates cell migration as well as cytoskeletal remodelling (80). Additionally, our group recently revealed that PANX1 co-immunoprecipitates with IQ motif-containing GTPase-activating protein 1 (IQGAP1), a multidomain scaffold protein that coordinates diverse cellular processes including cytoskeletal regulation partly through its direct interaction with F-actin which is known to directly interact with the C-terminus of PANX1 (36,57,106). It is notable that actin polymerization has been shown to play a critical role throughout cell division through its involvement in centrosome separation which we speculate could contribute to the increased proportion of cells stalling at G2/M in our adherent culture conditions when PANX1 is ablated (107,108). Altogether, our findings provide support for PANX1 functioning as a scaffold protein to support the structural assembly and integrity of melanoma spheroids.

Since our findings highlight PANX1 as a key mediator of melanoma cell adhesion, we sought to investigate whether supplementing extracellular matrix interactions with Matrigel would be sufficient to support the structural integrity of melanoma tumours in a chick chorioallantoic membrane xenograft model. However, despite the addition of Matrigel, we found that PANX1-deficient tumours exhibited reduced *en face* surface area coupled with decreased cell viability which is consistent with our previous studies in mouse melanomas, where the shRNA knockdown of PANX1 reduced tumour weight and occurrence of spontaneous metastasis (30). Taken together, our findings highlight a role for PANX1 in modulating the spatial organization and viability of melanoma tumours *ex vivo*. Although, we did not observe any differences in tumour characteristics upon treatment with spironolactone, we acknowledge this could be a limitation of the drug delivery method or short duration of tumour growth. To overcome this limitation, a syngeneic mouse model may be better suited to assess the potential tumour suppressive effects of spironolactone. Moreover, this would support the consideration of immune contributions which are key for the treatment of metastatic melanoma as immunotherapies are the current standard of care (7–9). Recently, our group revealed that the deletion of *PANX1* resulted in increased CD8^+^ cytotoxic T cell infiltration in tumours of an inducible melanoma model (29). Similarly, in colon cancer spironolactone was found to support immune surveillance by increasing the levels of Natural killer group 2D ligands (NKG2DLs) which interact with NKG2D receptors expressed on natural killer and cytotoxic T cells (109). Thus, future studies could assess whether spironolactone can synergize with PANX1 targeting strategies to improve the efficacy of immunotherapies.

Overall, this investigation highlights PANX1 as a potential therapeutic target for melanoma since the ablation of the *PANX1* gene and/or inhibition of its channel function altered cell cycle progression and the structural architecture of melanoma tumours. We demonstrate that the deletion of *PANX1* compromised spheroid compaction, reduced spheroid adhesion, as well as reduced viability and radial tumour growth in a chick chorioallantoic membrane xenograft model. Based on our bulk RNA sequencing and proteomic analysis, we speculate that PANX1 may function as a scaffold protein to support cell-cell and cell-ECM interactions that are critical for the 3D assembly and stability of melanoma. In addition, our data revealed that the deletion of PANX1 sensitizes melanoma cells to the pleotropic effects of spironolactone which could be exploited as an effective strategy to induce synthetic lethality in melanoma.

## ACKNOWLEDGEMENTS

Research reported in this publication was supported by funding from Schulich School of Medicine and Dentistry, Canada Foundation for Innovation, Ontario Research Fund, and NSERC Research Tools and Instruments Awards to Western University (London, Canada) through use of the Biomolecular Core (BioCORE) Facility.

## AUTHOR CONTRIBUTIONS

Conceptualization: S.E.L., D.W.L., S.P.

Data Curation: C.A.B.

Formal Analysis: S.E.L., B.L.O’D., W.J.S., M.S.H.

Funding Acquisition: B.E.I., J.R., D.W.L., S.P.

Investigation: S.E.L., D.J., K.B., B.L.O’D., W.J.S., M.S.H., D.S., C.A.B., J.J.K., Y.X.

Methodology: S.E.L., D.J., B.L.O’D., W.J.S., M.S.H., D.S., C.A.B.

Project Administration: S.E.L., D.J., S.P.

Resources: D.J., C.A.B., J.J.K., B.E.I., J.R., D.W.L., S.P.

Supervision: C.A.B., J.J.K., B.E.I., J.R., D.W.L., S.P.

Visualization: S.E.L., B.L.O’D., W.J.S., M.S.H.

Writing – original draft: S.E.L., B.L.O’D., C.A.B.

Writing – review and editing: all authors.

## SUPPLEMENTARY MATERIAL

**Supplementary Figure 1: The genetic ablation of PANX1 does not alter melanoma cell growth.** Immunoblotting using an antibody targeting the **(A)** amino- (PANX1 Cell Sig.) or **(B)** carboxy-terminus (PANX1 (CT-412)) of PANX1 confirms the successful generation of 6 distinct scrambled control (SC) (N=6, n=3) and 12 PANX1 knockout (KO) (N=12, n=3) A375-P clones using CRISPR-Cas9 gene editing. **(C)** Representative images of SC (N=6, n=1), and PANX1 KO A375-P (N=12, n=1), cells labelled with PANX1 (CT-412) (green). Nuclei were stained with Hoechst 33342 (blue). Scale bar = 10 µm. Clones highlighted in pink were selected as representative clones for downstream analysis. **(D)** The growth of SC (N=6, n =3-4) and PANX1 KO (N=12, n =3-4) A375-P clones were assessed over a period of 96 hours revealing no differences in cell growth when PANX1 was ablated from A375-P cells. **(E)** Average area under the curve (AUC) quantification for each SC and PANX1 KO clone (demarcated by shape) indicates variability in the AUC of distinct PANX1 clones but no overall difference in cell growth upon *PANX1* deletion. Blue and green data points denote the mean AUC for each independent CRISPR clone while the grey data points depict each independent biological replicate. Two-Tailed Unpaired T-Test, error bars are mean ± s.e.m.

**Supplementary Figure 2: The transcriptional profile of A375-P cells clusters according to the mechanism by which PANX1 is targeted.** SC (N=4) and PANX1 KO (N=4) A375-P cells were treated with 10 μM spironolactone (SPIR) or vehicle control (Veh.) for 96 hours and subsequently isolated for bulk RNA sequencing analysis. Principal component analysis (PCA) indicates clustering of samples according to each experimental condition **(A)** PANX1 ablation, **(B)** SPIR treatment in PANX1 expressing SC A375-P cells, or **(C)** SPIR treatment in PANX1 KO A375-P cells.

**Supplementary Figure 3: Cell cycle pathways are negatively enriched with the deletion of *PANX1* or treatment with spironolactone.** Gene set enrichment analysis (GSEA) of bulk RNA sequencing from SC (N=4) and PANX1 KO A375-P (N=4) cells indicates that Hallmark **(A)** E2F Target and **(B)** G2/M Checkpoint pathways are negatively enriched following the genetic ablation of *PANX1* and/or treatment with spironolactone (SPIR). Veh., vehicle.

**Supplementary Figure 4: The genetic ablation or pharmacological inhibition of PANX1 does not alter molecular indicators of endoplasmic reticulum stress, autophagic flux, or apoptosis.** Representative immunoblot of SC (N=4, n=4) and PANX1 KO A375-P cells (N=4, n=4) treated with 10 μM spironolactone (SPIR), 1 mM probenecid (PBN) or vehicle (Veh.) control probed with (A) C/EBP homologous protein (CHOP), (D) binding immunoglobulin protein (BiP) (G) Microtubule-associated proteins 1A/1B light chain 3B (LC3) and (J) cleaved caspase 3 (cCAS3). As a positive control for ER stress, SC A375-P cells were treated with tunicamycin (TM) and for apoptosis SC A375-P cells were treated with Staurosporine (Stauro.). Quantification revealed that the ablation and inhibition of PANX1 does not elevate the endoplasmic reticulum stress indicators (B-C) CHOP or (E-F) BiP, enhance autophagic flux (H-I) LC3-II/I, or promote apoptotic cell death (K-L) cCAS3. Two-way ANOVA with Tukey post-test.

**Supplementary Figure 5: PANX1 inhibitors do not alter the levels or cell surface localization of PANX1 in A375-P cells.** Representative immunoblot of PANX1 levels in SC A375-P clones treated with **(A)** 10 μM spironolactone (SPIR) or **(C)** 1 mM probenecid (PBN) with respective vehicle control (Veh.). **(B, D)** Quantification revealed that the inhibition of PANX1 channel function does not alter the absolute levels of PANX1 protein in SC A375-P cells (N=4, n=4). Two-Tailed Paired T-Test. Blue data points denote the mean for each independent CRISPR clone while the grey data points depict each independent biological replicate. There were no overt changes to subcellular distribution of PANX1 signal (green; PANX1(CT-412)) in pooled SC (n=3) A375-P cells following **(E)** 10 μM SPIR or **(F)** 1 mM PBN treatment relative to vehicle treated SC cells. Nuclei were stained with Hoechst 33342 (blue). Scale bar = 10 µm.

**Supplementary Figure 6: Inhibiting or deleting PANX1 does not alter the migration of adherent A375-P cells in scratch wounds. (A;** SPIR, **E;** PBN) Schematic illustrating the experimental timeline of PANX1 inhibition with 10 μM spironolactone (SPIR), 1 mM probenecid (PBN), or the corresponding vehicle (Veh.) control. **(B;** SPIR, **F;** PBN) Scratch wound closure was assessed over a 48-hour period in SC and PANX1 KO (n=3) A375-P cells and no differences in cell migration were observed. **(C;** SPIR, **G;** PBN) Wound closure was significantly reduced when PANX1 KO A375-P cells (n=3) were challenged with PBN but no changes in cell migration were observed with SPIR treatment. Blue and green data points denote the mean for each biological replicate while the grey data points depict technical replicates (4-6 scratched wells per experimental condition). Two-Tailed Paired T-Test. **(D;** SPIR, **H;** PBN) Representative phase contrast images of SC and PANX1 KO A375-P cells 48 hours following wounding. White lines depict the initial scratch would area. Scale bar = 400 µm.

**Supplementary Figure 7: *PANX1* and *ERCC3* transcript expression are positively correlated.** Gene Expression Profiling Interactive Analysis (GEPIA) of The Cancer Genome Atlas Skin Cutaneous Melanoma patient cohort reveals that the expression of PANX*1* and *ERCC3* mRNA are positively correlated.

