## Supplementary Figures for "Pannexin 1 Ablation Disrupts Cell Cycle Progression and Compromises the Structure of Melanoma Tumours"

### Supplementary Figure 1

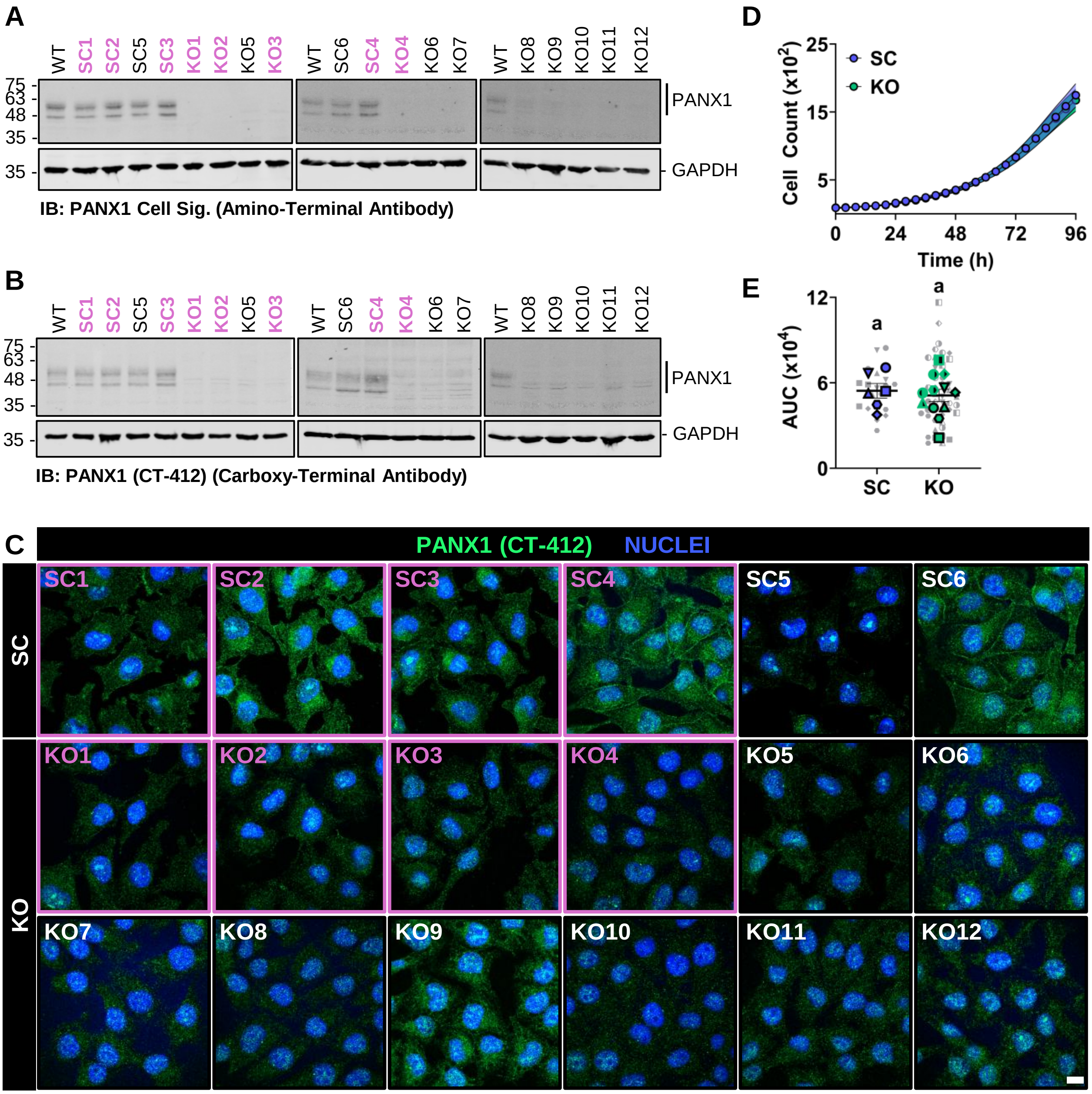

Supplementary Figure 2

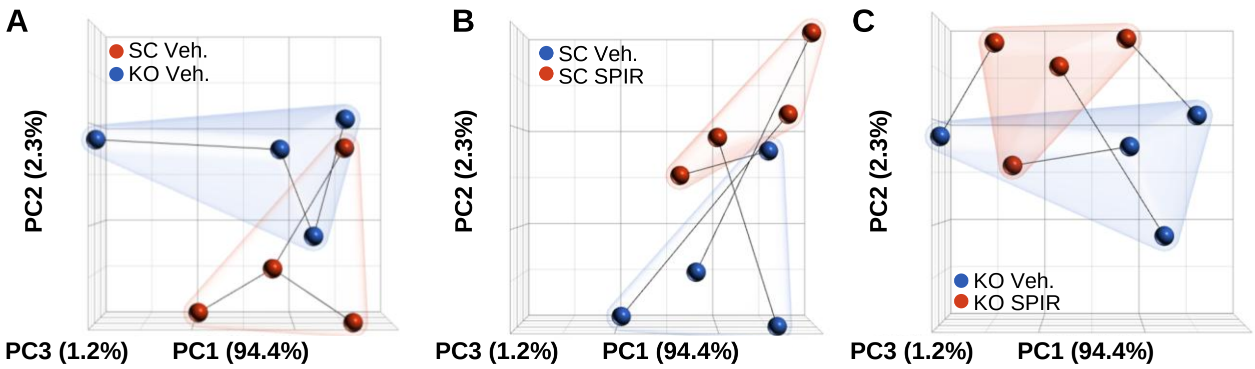

### Supplementary Figure 3

**A**

GSEA: HALLMARK E2F TARGETS

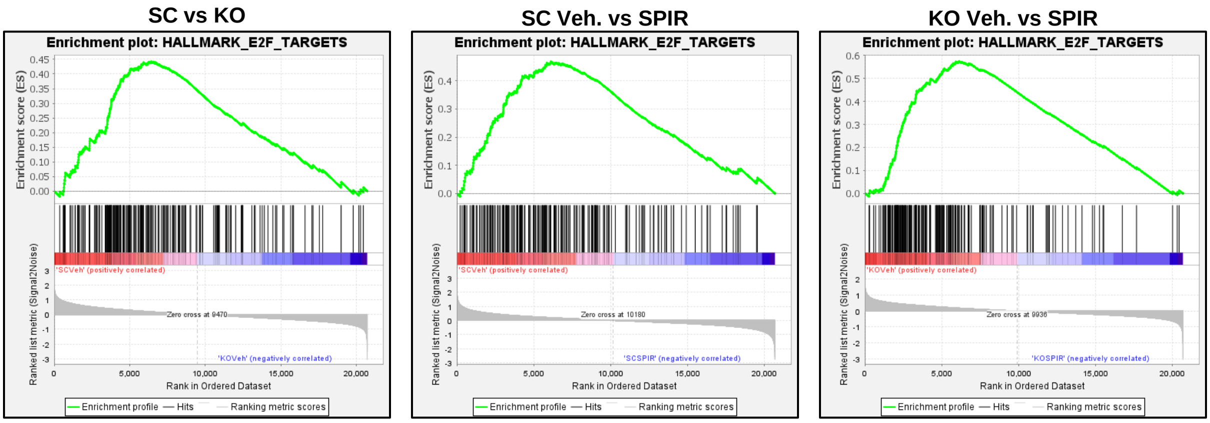


**B**

GSEA: HALLMARK G2M CHECKPOINT

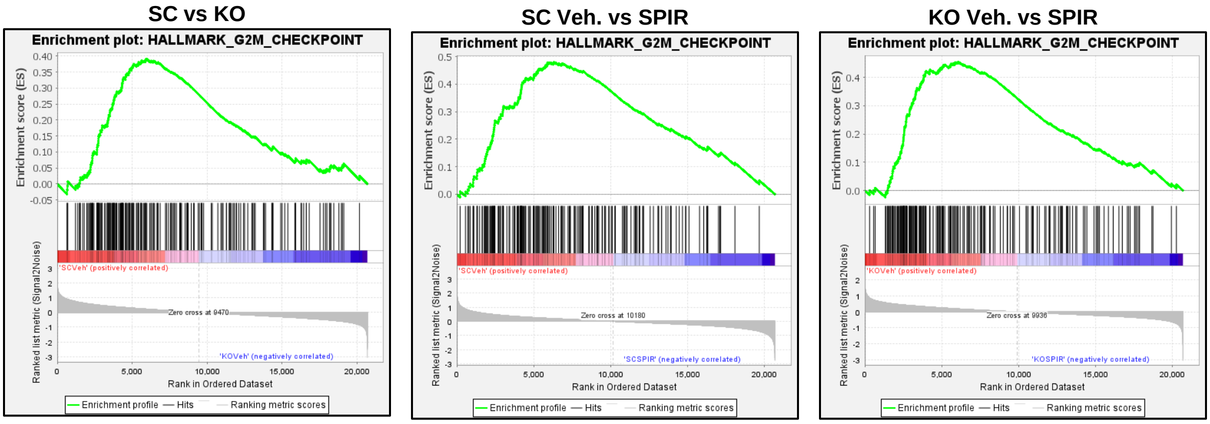

### Supplementary Figure 4

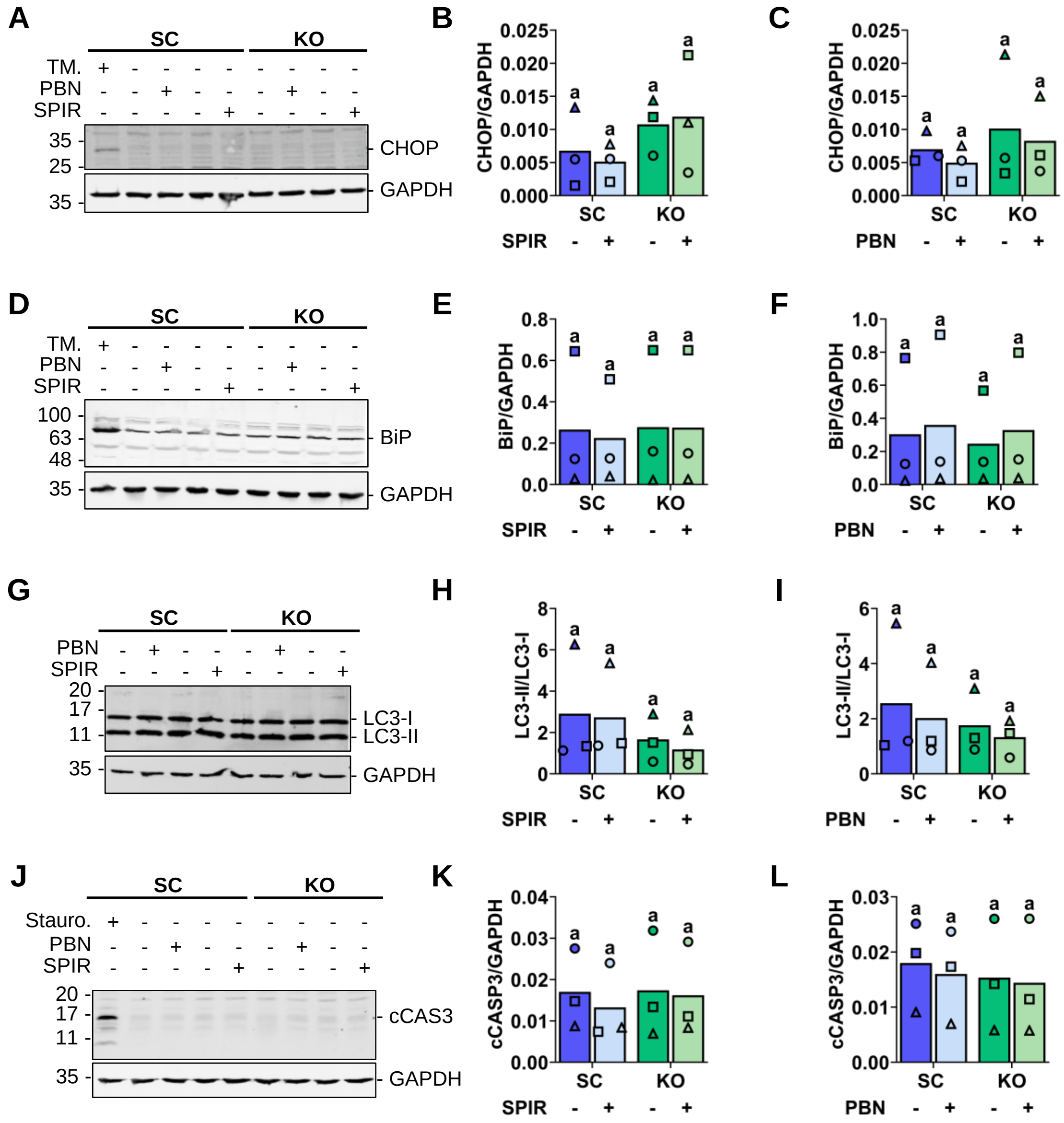

#### Supplementary Figure 5

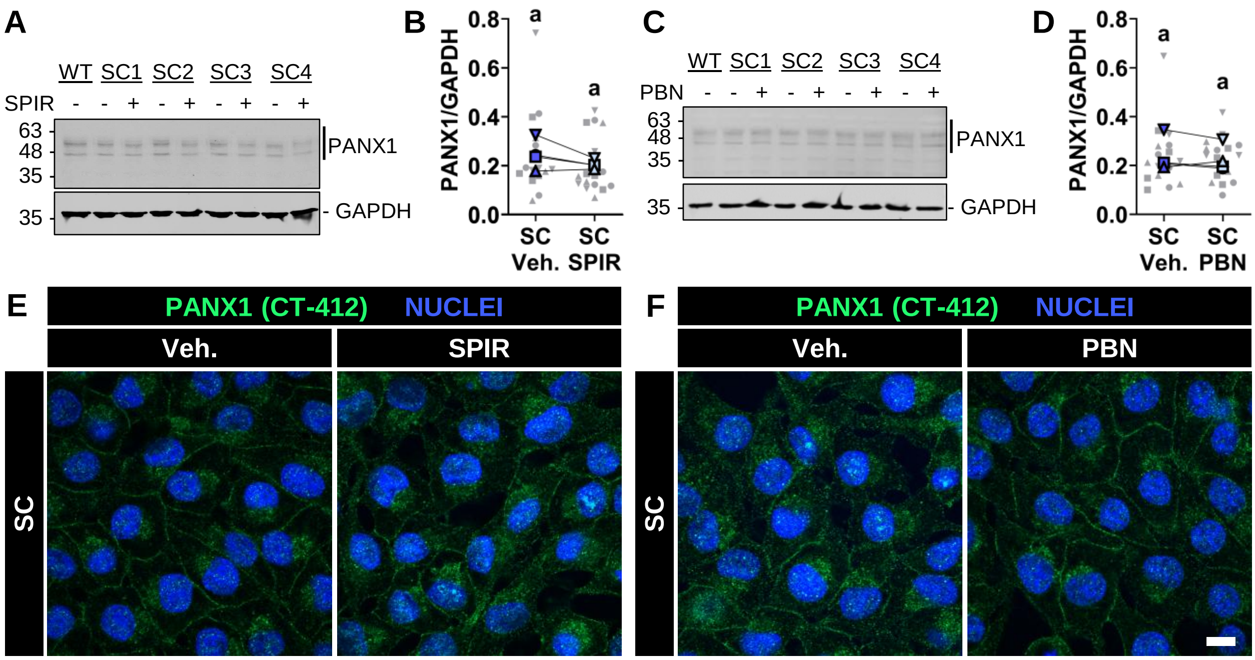

### Supplementary Figure 6

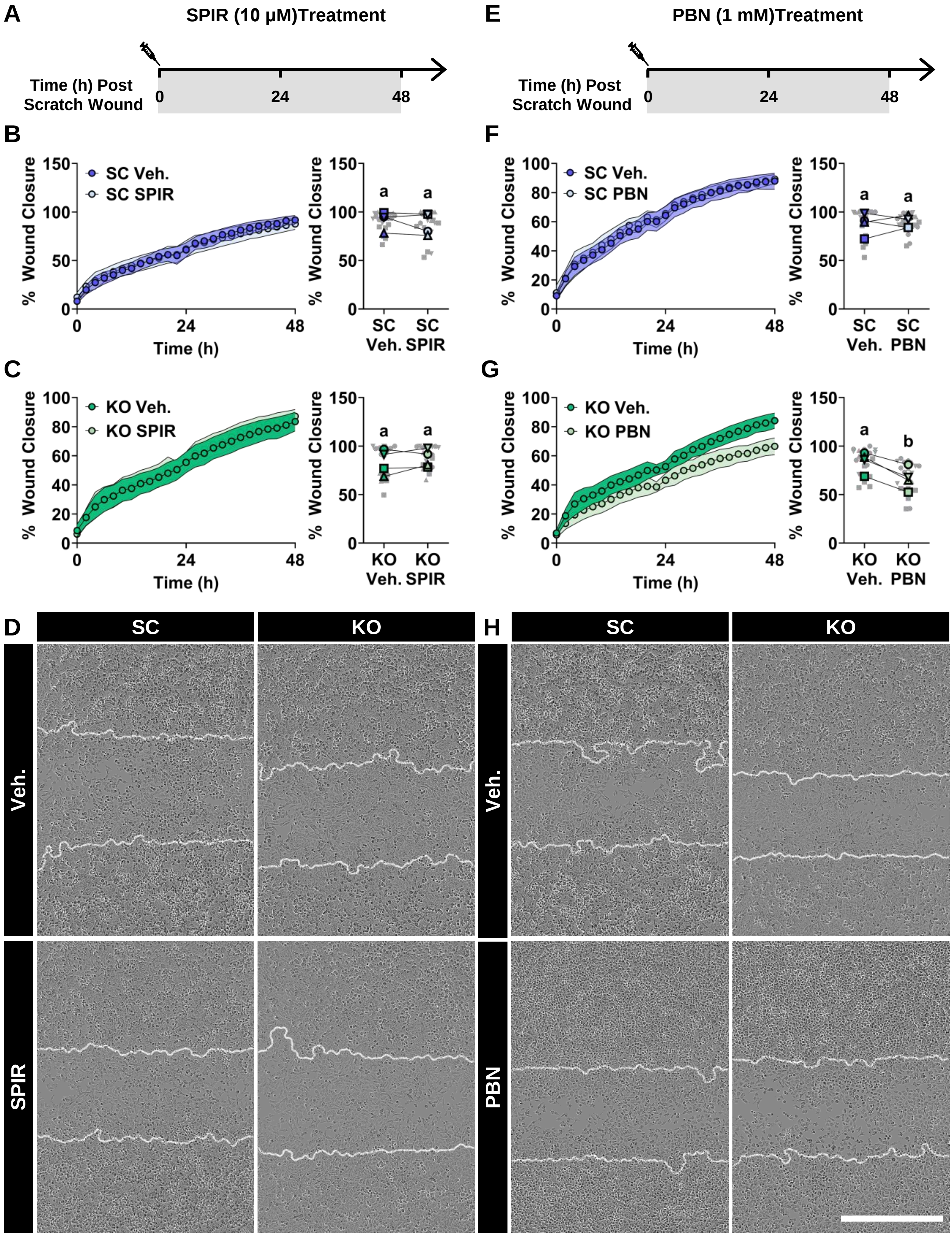

**Supplementary Figure 7**

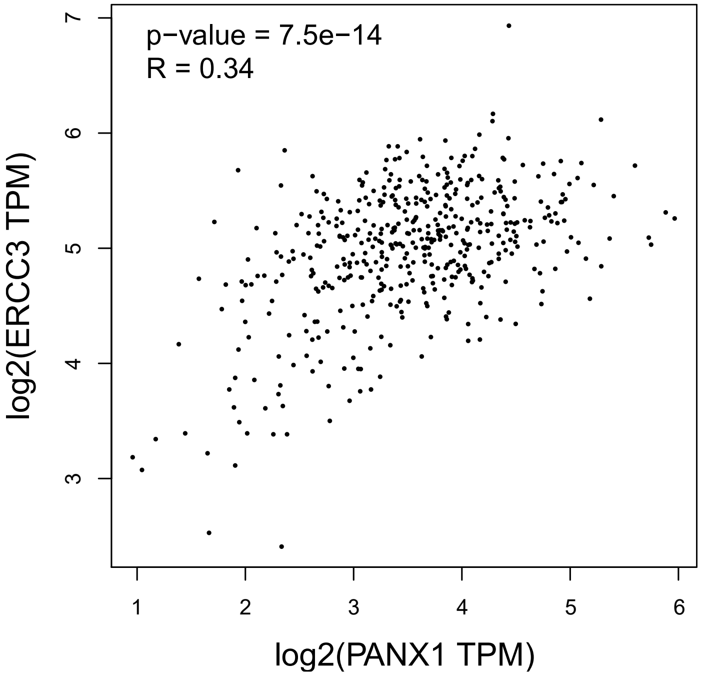
